# Assessing genome-wide dynamic changes in enhancer activity during early mESC differentiation by FAIRE-STARR-seq

**DOI:** 10.1101/2021.06.04.446899

**Authors:** Laura V. Glaser, Mara Steiger, Alisa Fuchs, Alena van Bömmel, Edda Einfeldt, Ho-Ryun Chung, Martin Vingron, Sebastiaan H. Meijsing

## Abstract

Embryonic stem cells (ESCs) can differentiate into any given cell type and therefore represent a versatile model to study the link between gene regulation and differentiation. To quantitatively assess the dynamics of enhancer activity during the early stages of murine ESC differentiation, we analyzed accessible genomic regions using STARR-seq, a massively parallel reporter assay. This resulted in a genome-wide quantitative map of active mESC enhancers, in pluripotency and during the early stages of differentiation. We find that only a minority of accessible regions is active and that such regions are enriched near promoters, characterized by specific chromatin marks, enriched for distinct sequence motifs, and modeling shows that active regions can be predicted from sequence alone. Regions that change their activity upon retinoic acid-induced differentiation are more prevalent at distal intergenic regions when compared to constitutively active enhancers. Further, analysis of differentially active enhancers verified the contribution of individual TF motifs toward activity and inducibility as well as their role in regulating endogenous genes. Notably, the activity of retinoic acid receptor alpha (RARα) occupied regions can either increase or decrease upon the addition of its ligand, retinoic acid, with the direction of the change correlating with spacing and orientation of the RARα consensus motif and the co-occurrence of additional sequence motifs. Together, our genome-wide enhancer activity map elucidates features associated with enhancer activity levels, identifies regulatory regions disregarded by computational prediction tools, and provides a resource for future studies into regulatory elements in mESCs.

## Introduction

Gene expression in eukaryotic cells is a tightly regulated process which is a prerequisite for cellular identity as well as any important cellular process. Regulation of transcription is controlled by transcription factors (TF) and the regulatory genomic elements (enhancers, promoters) they target (Ernst *et al*., 2011; Dunham *et al*., 2012). The selective and combinatorial activation of enhancers in a spatiotemporal manner allows for the complexity of higher eukaryotic organisms, which consist of a large number of different highly specialized cells although they all possess the same genome (Bulger and Groudine, 2011; Buecker and Wysocka, 2012). Traditionally, enhancers are defined as the genomic elements that can control the activity of promoters whereas promoters are the regions where transcription of genes is initiated. Further, promoters and enhancer regions can be distinguished and predicted based on distinct patterns of histone modifications (HMs) (Heintzman *et al*., 2007). However, recent research indicates that the function of enhancers and promoters may not always be distinct as studies have demonstrated that promoters can act as enhancers for other genes (Dao *et al*., 2017; Diao *et al*., 2017; Dao and Spicuglia, 2018) and enhancers frequently give rise to transcripts (de Santa *et al*., 2010), a feature traditionally associated with promoter function. The assignment of enhancers to their target promoters is an important step in elucidating gene regulation and has been addressed in recent years with rapidly evolving high-throughput chromatin interaction assays (Belton *et al*., 2012; Li *et al*., 2014; Mumbach *et al*., 2016). However, the functional relevance of identified enhancer-promoter pairs was mainly investigated for individual genes or loci (Sanjana, Shalem and Zhang, 2014; Canver *et al*., 2015; Diao *et al*., 2016; Korkmaz *et al*., 2016; Rajagopal *et al*., 2016; Gasperini *et al*., 2017; Klann *et al*., 2017) and remains a largely unsolved problem at the genome-wide level.

Further, gene expression is influenced by chromatin accessibility of regulatory elements and correlates with specific post translational HMs (Klemm, Shipony and Greenleaf, 2019). In eukaryotes, DNA is wrapped around a histone octamer to form nucleosomes, which are then organized into higher order chromatin. Chemical modifications of the histones tails demark promoters or enhancers and correlate with their transcriptional activity (reviewed in Buecker and Wysocka, 2012; Calo and Wysocka, 2013). The identification and prediction of enhancers is often based on indirect measures of activity, such as correlating HMs and chromatin accessibility (Ernst *et al*., 2011; Rajagopal *et al*., 2013; Zhu *et al*., 2013; Ernst and Kellis, 2017; Ramisch *et al*., 2019). Notably, some enhancer prediction tools discard promoter regions as potential enhancers even though there is evidence showing that promoters can act as enhancers of other genes. Moreover, enhancer prediction based on these marks gives rise to myriads of putative enhancers but doesn’t provide quantitative information regarding their activity. This is of particular interest, since gene expression is not subject to an on/off switch type of regulation, but rather the result of a complex interplay between multiple enhancers, TFs, and coactivators which can fine-tune gene expression levels to meet the cell’s current needs. Consequently, it remains largely unclear which of the thousands of predicted enhancers are actually functional, how enhancer usage changes during differentiation and what features are conferring distinct activity levels.

Embryonic stem cells (ESCs) are characterized by their ability to differentiate into any given cell type and therefore represent a versatile system to study the link between gene regulation, differentiation and cellular identity (Silva and Smith, 2008; Young, 2011). Murine ESCs (mESCs) in the pluripotent state exhibit relatively permissive chromatin, with many accessible regions which are thought to comprise active mESC enhancers but also primed enhancers that can be activated at later stages during differentiation (Buecker *et al*., 2014; Wu *et al*., 2016). The expression of genes in mESCs is also controlled by transposable elements, for example from the ERVK family, that can act as enhancers that control the expression of associated genes (Sundaram *et al*., 2017; Todd *et al*., 2019; Hermant and Torres-Padilla, 2021). The pluripotency of mESCs and their ability to self-renew critically depend on the actions of specific TFs including OCT4 and SOX2, NANOG, KLF4, and ESRRB (Chambers and Tomlinson, 2009; Young, 2011). All these TFs can bind and activate promoters as well as enhancers of pluripotency-associated genes in ESCs (Buecker *et al*., 2014). mESCs can be cultivated in the pluripotent state when leukemia inhibitory factor (LIF) is added to the media to activate the STAT3 pathway (Niwa *et al*., 1998), which in turn promotes c-MYC expression and transcriptional programs important for self-renewal (Cartwright *et al*., 2005).

Differentiation of ESCs can be used to study the molecular mechanisms that underly cellular commitment decisions with potential therapeutic relevance. Many differentiation protocols for diverse cell types have been established. However, many of these protocols suffer from low differentiation efficiencies which limits their applicability. A highly efficient, yet simple, protocol to induce cellular differentiation is to treat mESCs with all-trans retinoic acid (RA) (Gudas and Wagner, 2011). Treatment with RA induces exit from pluripotency, marks a phase of increased susceptibility to lineage-defining signals (Semrau *et al*., 2017) and ultimately pushes mESCs towards the neuronal lineage (Janesick, Wu and Blumberg, 2015). RA is the ligand of retinoic acid receptors (RARs), which together with retinoid X receptors (RXRs) bind to genomic response elements and drive expression of differentiation-associated genes but also repression of genes involved in pluripotency (Chatagnon *et al*., 2015). Among the targets of RA-induced differentiation are the well-studied *Hoxa* genes, which are coding for TFs that play a pivotal role in development and body axis formation (Neijts and Deschamps, 2017).

In recent years, several studies applying massive parallel reporter assays based on STARR-seq (Arnold *et al*., 2013) have been conducted to assess enhancer function of candidate regions in different species and cell types (Shlyueva *et al*., 2014; Vanhille *et al*., 2015; Dao *et al*., 2017; Barakat *et al*., 2018; Schöne *et al*., 2018; Wang *et al*., 2018; Chaudhri *et al*., 2020; Peng *et al*., 2020). Here, we developed a modified quantitative STARR-seq protocol focusing on accessible chromatin, thereby including promoter and enhancer regions, as candidate enhancers. This allowed us not only to identify active enhancers genome-wide in mESCs, but also to quantify enhancer activity and thus to identify features, such as sequence motifs and their quantities, that correlate with enhancer activity. Moreover, we used our quantitative approach to study enhancers upon differentiation to identify those that change their activity during the early stages of differentiation. Additionally, we intersected RARα binding with RA-induced changes in enhancer activity to identify “functional” RARα binding sites. This resulted in the identification of sequence features associated with RARα binding events with distinct changes in enhancer activity.

Overall, our studies using the FAIRE-STARR-seq assay provide a genome-wide resource for enhancer activity levels for mESCs in the pluripotent state and after induced differentiation and uncovers features that are important for enhancer activity and consequently might play a role in modulating expression levels of associated genes.

## Material and methods

### mESC culture and differentiation

E14 mESCs were cultured under feeder-free conditions and routinely passaged every two days in ES-medium: Glasgow Minimum Essential Medium (Sigma-Aldrich) supplemented with 17% FBS (Hyclone™, SV30160.03, GE Healthcare), 2 mM GlutaMAX™ (Gibco), 100 U/ml Penicillin-Streptomycin (Gibco), 1x MEM Non-Essential Amino Acid Solution (Gibco), 1 mM Sodium Pyruvate (Gibco), 0.5 mM 2-Mercaptoethanol (Gibco), and recombinantly expressed leukemia inhibitory factor (LIF). To exit from pluripotency and induce differentiation, LIF was withdrawn and retinoic acid (RA, Sigma, R2625) was added to the medium to a final concentration of 1 µM. For all experiments, 4 h prior to harvest, cell culture medium was removed, cells washed twice with PBS and fresh medium containing either LIF or RA was added.

### FAIRE-STARR-seq

As input material for the reporter screen, accessible chromatin from E14 mESCs treated with RA (Sigma, #R2625) for 4 h was isolated by formaldehyde-assisted isolation of regulatory elements (FAIRE, Giresi *et al*., 2007) and subsequently cloned into the STARR-seq screening vector (Addgene #71509) following the protocol described in Arnold *et al*., 2013. To asses enhancer activity, E14 cells were transfected with this plasmid library using a Nucleofector™ 2b device using the Mouse ES Cell Nucleofector Kit (Lonza, VAPH-1001). For each of the three biological replicates, four individual transfections, each with 5 µg plasmid library and 5×10^6^ cells, were performed. The medium was changed 12 h after transfection and to half of the cells either LIF or 1 µM RA was added. After an additional 4 h of incubation, samples were pooled and RNA was isolated using the Rneasy Midi kit (Qiagen). Poly adenylated RNA was enriched using Dynabeads™ Oligo(dT)_25_ (Invitrogen), residual DNA was digested using Turbo DNase (Invitrogen), and finally RNA was cleaned-up with Agencourt^®^ RNAClean^®^ XP beads (Beckman Coulter). cDNA was synthesized using SuperScript™ III Reverse Transcriptase (Invitrogen) according to the manufacturer’s protocol, applying a reporter transcript-specific primer. This primer contains the sequence of the Illumina PCR Primer 2.0 as overhang as well as eight random nucleotides that serve as unique-molecular identifiers (UMI) for each cDNA molecule (CAAGCAGAAGACGGCATACGAGAT[N]_8_ GTGACTGGAGTTCAGACGTGTGCTCTTCCGATCT). cDNA was further amplified as described in Arnold *et al*., 2013, using adjusted reporter-specific primers based on Illumina’s TruSeq dual index system (universal: CAAGCAGAAGACGGCATACGA, sample specific: AATGATACGGCGACCACCGAGATCTACAC[barcode, n=6]ACACTCTTTCCCTACACGACGCTC).

As input control for FAIRE-STARR-seq, the input plasmid library was sequenced as well. To this end, the plasmid library was used for a pseudo “cDNA synthesis”, using the random-UMI primer and the KAPA HiFi HotStart ReadyMix (Roche) for 4 cycles with a prolonged synthesis step (70 sec) to individually label input fragments. In a second step, this input library was amplified with Illumina’s TruSeq dual index based universal and barcoded primers, as done for the FAIRE-STARR-seq libraries, using the KAPA HiFi HotStart ReadyMix (Roche) for 12 PCR cycles.

### STARR-qPCR

Putative enhancer sequences (Table S1) were amplified by nested PCR from genomic DNA derived from E14 cells using standard PCR procedures. Primers (Table S1) were designed to generate the same overhangs as used for Illumina sequencing. The negative (nc1 and nc2, GR responsive elements) and positive (CMV enhancer) control regions as well as RARα motif variants were ordered as gBlocks (IDT) and are listed in Table S1. DNA fragments were subsequently cloned into the STARR-seq screening vector (pSTARR-seq_human, Addgene plasmid #71509) using the In-Fusion^®^ HD Cloning Kit (Takara/Clonetech). For transfection of reporter plasmids, E14 mouse ESCs were plated at a density of 1.4 × 10^4^ cells/cm^2^ of a 24 well plate with ES medium supplemented with 17% FBS and LIF. The next day, cells were washed with PBS and fresh medium was added. Subsequently, cells were transfected with individual reporter plasmid using Lipofectamin 2000 (Invitrogen) according to the manufacturer’s instructions. 20 h after transfection, cells were washed twice with PBS and fresh ES medium, containing LIF, 1 µM RA or no additional reagent, was added. After another 4 h of incubation, cells were harvested, RNA extracted (RNeasy Mini Kit, Qiagen), followed by cDNA synthesis (PrimeScript RT Reagent Kit, Takara, using oligodT and random hexamer primers). Reporter transcript levels were quantified by qPCR with primers specific for GFP and normalized to the expression of two housekeeping genes (*Rpl19* and *Actb*). Primers are listed in Table S1.

### ATAC-, ChIP-, and RNA-seq experiments

ATAC-, HM ChIP-, and RNA-seq experiments from our laboratory have been published previously (Ramisch *et al*., 2019). RARα ChIP was performed for this study. In short:

#### ATAC-seq

75,000 low passage (< 10) E14 cells were cultivated for 48 h in ES medium prior to subjecting them to an improved ATAC-seq protocol as described in Corces *et al*. (2017). The resulting transposase-fragmented and PCR-amplified DNA was cleaned up using AMPure XP beads (Agencourt). High-throughput sequencing was performed generating approx. 50 million 50 bp paired-end reads per sample using the HiSeq 4000 (Illumina) device

#### ChIP-seq

For ChIP experiments, E14 cells were washed once with PBS, treated with trypsin (Sigma, T4049) for 5 min and gently but thoroughly resuspended in ES medium to generate single cell suspensions. Cells were diluted to 20×10^6^ cells/20 ml medium and crosslinked by adding formaldehyde (1% v/v) for 5 min under gentle rotation. The reaction was quenched by adding 125 mM Glycine for an additional 5 min, then cells were washed three times with PBS, snap frozen in liquid nitrogen, and stored at -80°C.

<u>HM ChIP</u> experiments were preformed according to the standard BLUEPRINT protocol (www.blueprint-epigenome.eu): Cells were resuspended in shearing buffer (20 mM Tris pH 7.5, 150 mM NaCl, 2 mM EDTA, 1% Triton X-100, 0.1% SDS) supplemented with Complete Protease Inhibitor Cocktail (PIC) EDTA-free (Roche, 11873580001) and sheared on a Bioruptor Pico device for 25-35 cycles. For each ChIP, 1 µg antibody (listed in Table S2) was used. Automatic ChIP was performed using the SX-8G Compact IP-Star liquid handler (Diagenode) in combination with Auto Histone ChIP kits (Diagenode, C01010022). Using the pre-programmed method ‘indirect ChIP’, ChIP reactions were carried out in a final volume of 200 μl for 10 h followed by 5 h incubation with protein A magnetic beads and 5 min washes at 4°C. After the ChIP, eluates were recovered, RNase A-treated, de-crosslinked overnight at 65°C and treated with Proteinase K for 4 h at 55°C. The recovered DNA was purified using the ChIP DNA Clean & Concentrator Kit (Zymo research, D5205). Sequencing libraries were prepared using the NEBNext Ultra DNA Library Prep kit (NEB, E7370) according to manufacturer’s instructions and submitted for paired-end Illumina sequencing on the HiSeq 2500.

The <u>RARα ChIP</u> was performed as described elsewhere (Glaser *et al*., 2017), with the following modifications: Cells were cross-linked for 5 min with 1% formaldehyde and a mild sonication buffer was used (20 mM Tris-HCl pH 8.0, 2 mM EDTA pH 8.0, 1% Triton X-100, 150 mM NaCl, 0.1% SDS, 1x PIC). Prior to sonication, nuclei were incubated for 20 min on ice and homogenized ten times by a 27G needle. Per ChIP 4 µl RARα antibody (serum, Diagenode C15310155) or 2 µg IgG control (Diagenode C15410296) was used. Sequencing libraries for RARα ChIP and Input fragments were prepared using the KAPA Hyper Prep Kit (Roche) and submitted for paired-end Illumina sequencing on the NovaSeq 6000 generating 50 bp reads.

#### RNA-seq

2×10^5^ low passage (< 10) E14 cells were plated per 10 cm dish and cultivated for 48 h in regular ES medium. Next, medium was exchanged for fresh ES medium containing either LIF or 1 µM retinoic acid (Sigma, R2625). After 4 h, cells were harvested and RNA extracted using the RNeasy Mini Kit (Qiagen) according to the manufacturer’s instructions. The experiment was performed in biological triplicates. Sequencing libraries were generated using the TruSeq^®^ Stranded mRNA Kit (Illumina) and high-throughput sequencing was performed on a HiSeq 2500 (Illumina) device generating approx. 100 million 50 bp paired-end reads per sample.

### Generation of clonal cell lines with CRISPR/Cas9-mediated genomic deletions and mutations

sgRNAs targeting regions of interest were designed using the CRISPOR tool (http://crispor.org/, Concordet and Haeussler, 2018), ordered as complementary DNA oligonucleotides (Sigma) with overhangs for BbsI, and cloned into the pSpCas9(BB)-2A-Puro (PX459) V2.0 plasmid (Addgene plasmid #62988) as described in Ran *et al*. (2013). All sgRNA sequences are listed in Table S1. To delete regions of interest, two million E14 cells were transfected with a pair of sgRNA plasmids (as indicated), 1 µg per plasmid, using a Nucleofector™ 2b device and the Mouse ES Cell Nucleofector Kit (Lonza, VAPH-1001) according to the manufacturer’s instructions and plated into two 10 cm dishes. 24 h after transfection, medium was exchanged for fresh ES medium, and after another 24 h the medium was exchanged for fresh ES medium containing 2.5 µg/ml Puromycin. The next day, medium was exchanged again for ES medium without selection. Subsequently, medium was exchanged every two days until round colonies formed (7-10 days post transfection). Colonies were picked by pipetting and individually transferred into 48 well plates. E14 clonal lines were expanded, genomic DNA was extracted (QIAamp DNA Mini Kit, Qiagen), and lines were genotyped using primers listed in Table S1 and Phusion High-Fidelity PCR Master Mix (with GC Buffer) (Thermo Scientific, F532). PCR products of candidate clonal lines showing predicted PCR band sizes in agarose gel electrophoresis, were send for validation by sanger sequencing (Eurofins). To probe for biallelic alterations, PCR products were cloned into the Zero Blunt™ vector (PCR Cloning Kit, Thermo, K270020), transformed into *E*.*coli*, four to eight individual bacterial colonies were picked, plasmid DNA isolated (QIAprep Spin Miniprep Kit, Qiagen) and send for Sanger sequencing. Genomic deletions and mutations of E14 clonal lines are listed in Table S3.

### RT-qPCR

RNA from E14 or clonal cell lines, treated as indicated, was extracted using the RNeasy Mini Kit (Qiagen) according to the manufacturer’s instructions including a DNase treatment. 1 µg total RNA was subjected to cDNA synthesis applying the ProtoScript^®^ First Strand cDNA Synthesis Kit (NEB, E6300S) with the included Oligo d(T)23 VN primer according to the manufacturer’s instructions. cDNA was diluted 1:12.5-1:20 prior to qPCR which was performed as described in Thormann *et al*. (2018).

### NGS data analyses

#### FAIRE-STARR-seq data analyses

FAIRE-STARR-seq libraries were sequenced with a HiSeq 2500 (Illumina) to generate 50 bp paired-end reads. Sequencing reads were aligned to the mouse genome (mm9) using Bowtie2 (Langmead and Salzberg, 2012)(-X 800 --fr --very-sensitive). UMI-tools (Smith, Heger and Sudbery, 2017) was used for UMI-aware removal of PCR duplicates. SAMtools (Li *et al*., 2009) was used to filter reads for proper pairs, alignment and quality scores (-h -b -f 3 -F 780 -q 5), to select reads mapping only to regular chromosomes (chr1-19, chrX and chrY), and to remove reads mapping to blacklisted regions (ENCFF547MET). UMI-aware deduplication of reads removed about 90% of obtained reads (Fig. S1A) and is aimed at retaining only true biological replicates resulting in an overall decrease in read-counts for individual fragments (Fig. S1B). Genome-wide correlation analyses of read distributions of individual FAIRE-STARR-seq samples showed higher correlation coefficients when UMI-aware removal of read duplicates was omitted. Fragments with extremely high read counts in only one replicate are prevalent without UMI-aware removal of duplicates, whereas these regions are absent after UMI-aware deduplication analysis (Fig. S1C) indicating that such regions are PCR amplification artefacts. Accessible regions covered by the input library were identified using MACS2 (Zhang *et al*., 2008)(-q 0.05 --keep-dup all --call-summits -bw 200). Significantly active enhancers, using the input library as control, were called using MACS2 (Zhang *et al*., 2008). The analysis was performed for each biological STARR-seq replicate individually as well as for the merged reads from all replicates. Finally, peaks were only counted as active STARR-seq enhancers when they were called for the merged reads and for at least two of three biological replicates and are covered by at least three individual fragments. Normalized STARR-seq signal for data visualization was generated using bamCoverage of the deepTools package (Ramírez *et al*., 2016) for the replicate-merged STARR-seq reads or the input library to normalize for genomic coverage and sequencing depth (-of bigwig -bs 10 -e --normalizeUsing RPGC --effectiveGenomeSize 2304947926 --pseudocount 1). Next, signal tracks were normalized to input library coverage using bigwigCompare (-of bigwig -bs 10 --operation subtract --pseudocount 1). Heatmaps which show STARR-seq signal distribution at selected regions were generated using computeMatrix (reference-point mode) and plotHeatmap tools of the deepTools package (Ramírez *et al*., 2016). Genomic distribution of FAIRE-STARR with respect to RefSeq genes was annotated with ChIPSeeker (Yu, Wang and He, 2015).

In order to score the FAIRE-STARR-seq enhancers, the computeMatrix tool of the deepTools package (Ramírez *et al*., 2016) was used, this time to obtain the average enhancer activity signal (input and read depth normalized tracks by bigwigCompare, see above (--operation log2)) over the size-scaled regions (scale-regions mode). Clustering of FAIRE-STARR enhancers by enrichment of HMs was performed using the computeMatrix tool (scale-region mode to average HM enrichment per region) and k-means clustering (k was estimated by the elbow method (total within-cluster sum of square)). Subsequently, distributions of HMs, TFs, accessibility by ATAC, promoter annotation (RefSeq), transcription (RNA-seq), and enhancer prediction probability by CRUP (Ramisch *et al*., 2019) were plotted for the clustered regions with computeMatrix (reference-point mode on summit of the clustered regions) and plotHeatmap (Ramírez *et al*., 2016).

#### Correlation analyses

Genome-wide correlation analyses for read distributions were performed using multiBamSummary (deepTools, Ramírez *et al*., 2016) and filtered reads. The genome was binned into 100 bp bins, fragments per bin were counted (bins -e -bs 100), the resulting table was analyzed in R (R Core Team, 2017) and pair-wise Pearson correlation coefficients and coefficients of determination were calculated.

#### ChIP-seq analyses

Paired-end ChIP-seq reads were mapped to the reference genome (mm9) using Bowtie2 (Langmead and Salzberg, 2012)(--sensitive), and if applicable, mapped reads from the same experiment but different sequencing runs were merged. SAMtools (Li *et al*., 2009) was used to filter for proper pairs, alignment and quality scores (-h -b -f 3 -F 780 -q 10), to select reads mapping only to regular chromosomes (chr1-19, chrX and chrY), and to remove reads mapping to blacklisted regions (ENCFF547MET). Input and sequencing depth normalized signal tracks were computed with bamCompare (-of bigwig --operation subtract -bs 25 --smoothLength 50 -e --normalizeUsing RPKM -- ignoreDuplicates) (Ramírez *et al*., 2016). Significant RARα binding sites over input sample were identified using MACS2 (Zhang *et al*., 2008). For RARα enhancer inducibility analysis, only RARα binding sites which overlap with the FAIRE-STARR input library (6,528 of 11,366 RARα sites) were included.

#### Reprocessing of deposited NGS data

If signal tracks were not available, NGS data for experiments listed in Table S2 were downloaded via fastq-dump, mapped to mm9 reference genome using Bowtie2 (Langmead and Salzberg, 2012)(-- sensitive), and if applicable, mapped reads from the same experiments but different sequencing runs were first merged and then filtered (-h -b -f 3 -F 780 -q 3) with SAMtools (Li *et al*., 2009). Signal tracks were computed with bamCoverage or bamCompare (-of bigwig (--operation subtract) -bs 25 -- smoothLength 50 -e --normalizeUsing RPKM --ignoreDuplicates) (Ramírez *et al*., 2016) depending on the availability of a control sample (indicated in Table S2). Reads mapping to blacklisted regions (Dunham *et al*., 2012) were excluded. For deposited signal tracks mapped to mm10 reference genome, lift-over to mm9 was performed using CrossMap (Zhao *et al*., 2014).

#### RNA-seq analysis

50 bp paired-end sequencing reads were aligned to the mouse genome (mm9) using STAR (Dobin *et al*., 2013)(version 2.5.3a) and ENSEMBL genes (NCBIM37) as annotation reference. SAMtools (Li *et al*., 2009) was used to filter reads for proper pairs, alignment and quality scores (-h -b -f 3 -F 780 -q 10), to select reads mapping only to regular chromosomes (chr1-19, chrX and chrY), and to remove reads mapping to blacklisted regions (ENCFF547MET). Fragments per gene were assessed using featureCounts (Liao, Smyth and Shi, 2014) and ENSEMBL gene annotation. To compare expression between different groups of genes of the same treatment, transcripts per million reads (TPM) were calculated and compared. Normalization of read coverage and differential gene expression analysis for different treatments were performed using DESeq2 and LCF shrinkage (Love, Huber and Anders, 2014). To compare and plot mean expression of genes between different treatments, ™M-normalized counts (Robinson and Oshlack, 2010) were calculated with the edgeR package (Robinson, McCarthy and Smyth, 2010). To generate signal tracks for plotting RPKM normalized read coverage at example loci or heatmaps, bamCoverage was used (-of bigwig -bs 10 -e --normalizeUsing RPKM)(Ramírez *et al*., 2016).

#### ATAC-seq analysis

50 bp paired-end sequencing reads were aligned to the mouse genome (mm9) and filtered as described for ChIP-seq analysis. Signal tracks for plotting normalized read coverage at example loci or heatmaps were generated applying bamCoverage (-of bigwig -bs 25 --smoothLength 50 -e -- normalizeUsing RPGC --effectiveGenomeSize 2304947926 --ignoreDuplicates)(Ramírez *et al*., 2016).

#### Motif enrichment analyses

To identify TF motifs enriched in sequences of interest, AME (McLeay and Bailey, 2010) was applied (--scoring avg --method fisher --hit-lo-fraction 0.25 --evalue-report-threshold 79 --control --shuffle--) using the JASPAR 2018 clustered vertebrate motif database (Khan *et al*., 2018) as input motifs. Results were analyzed in R (R Core Team, 2017), filtered by E-value thresholds as indicated, and plotted with the ggplot2 package (Wickham, 2009). The JASPAR 2018 vertebrate core motifs and their corresponding clusters are listed in Table S4. To investigate the enrichment of RARα::RXRα motifs with different spacer lengths and half-site orientations, the corresponding scoring matrices were created by combining the monomers of the RARα::RXRα consensus motif (MA0159.1) into direct, inverted, and everted repeats with zero to eight nucleotides spacing. For the spacers, a uniform nucleotide frequency distribution was inserted to generate maximal degeneracy.

Counting of enriched motifs per fragment was performed using the matrix-scan function of the pattern matching program from RSAT software suite (Turatsinze *et al*., 2008) with a first-order Markov model estimated from the input sequences as a background model and applying a p-value cut-off (0.002) to the predicted binding sites.

#### Heatmaps and anchor plots

Heatmaps and anchorplots depicting ChIP-, DNase-, ATAC-, or RNA-seq distribution or mean enrichment at selected genomic regions respectively, were generated using computeMatrix (reference-point mode) and subsequently plotHeatmap or plotProfile tools of the deepTools package (Ramírez *et al*., 2016). Sequencing depth and, if applicable and available, input normalized signal tracks were used.

#### Assignment of genes to enhancers and gene ontology analysis

To assign putative target genes to STARR enhancers we applied GREAT version 3.0.0 (McLean *et al*., 2010) using the whole genome (mm9) as background regions and for association setting “basal plus extension” with proximal: 5 kb upstream and 1 kb downstream, plus distal: up to 100 kb. The expression levels of assigned genes per enhancer group or cluster was plotted as TPM derived from RNA-seq. Additionally, GREAT performs a gene ontology analysis per analyzed enhancer group and provides enriched GO-terms and significance levels, which were analyzed and cutoffs determined in R (R Core Team, 2017) and subsequently plotted with the ggplot2 package (Wickham, 2009).

#### Classifier for enhancer and E-promoter prediction

##### Pre-processing and motif enrichment

As outlined in Fig. 4a, the 186,959 significantly enriched regions of the FAIRE-STARR input library were first divided into regions which do (16,769) or do not (170,190) overlap with ENSEMBL (NCBIM37) promoters, which were defined as regions of -500 bp to the TSS, and subsequently used to train an E-promoter and enhancer classifier, respectively. For each group, regions were ranked for their STARR activity (Fig. 4b and S4a) and the sequences of the highest and lowest ranking 10 or 1% for E-promoters or enhancers, respectively, were used for training of the classifier. The motifcounter tool (Kopp 2017) was used with default options to calculate sequence-wise motif enrichment of the 79 clustered motifs from JASPAR matrix clustering 2018 (Khan *et al*., 2018) using the union from both sets as background model. Since the width of highest and lowest STARR-scoring regions was significantly different (Wilcoxon p < 1e-50), region-width was included as a feature of the classifier. Negative log-transformed p-values of motif enrichment were generated and all variables were scaled such that they have the same mean and standard deviation, in order to allow for inferences about feature importance directly from regression model coefficients.

##### Fitting and evaluation of classifier

To differentiate between the highest and lowest ranking enhancers based on enrichment of the clustered TF motifs and motif width, a logistic regression model with elastic net regularization was built. The model combines ridge and lasso penalties to obtain shrunken and grouped coefficients, that prevent the regression model from overfitting (Zou and Hastie, 2005). For training and evaluation of the model, a nested cross-validation approach was performed, where the inner loop is used for the optimization of hyperparameter λ (regularization penalty) and the outer loop to assess the predictive performance on unseen data. Additionally, the second hyperparameter α was tested over a grid of various values to find the optimal mixing percentage of lasso and ridge regression. Since only marginal differences in performance were observed, a value of α = 0 corresponding to ridge regression was chosen to include enrichment of each of the clustered motifs in the classifier. Model performance for each of the outer cross-validation folds was assessed via the receiver operating characteristic (ROC) curve to derive a mean and standard deviation of the AU-ROC (area under the ROC curve). Preprocessing, training, and testing of the model were performed with R using the glmnet package (Friedman, Hastie and Tibshirani, 2010) for elastic-net regularized models.

## Results

### Generation of a quantitative enhancer-activity map of the mESC genome

To assess the enhancer activity of putative regulatory regions in mESCs, we performed a massively parallel enhancer reporter assay. To limit the complexity of the library, we prioritized regions that are likely to act as enhancers (Klemm, Shipony and Greenleaf, 2019) by focusing on accessible chromatin isolated by FAIRE (formaldehyde-assisted isolation of regulatory regions) as input material for our STARR-seq (self-transcribing active regulatory regions)(Arnold *et al*., 2013) library (Fig. 1a). Although this idea was new at the time of conception, a similar approach has by now also been described by others that isolated putative regulatory elements by either FAIRE (Chaudhri *et al*., 2020) or ATAC (Wang *et al*., 2018). To determine the complexity of the FAIRE fragments, we sequenced the plasmid input library (Fig. 1a) resulting in the identification of 4.4 million individual fragments, which cover about 186,000 significantly enriched open regions. As expected, the enrichment of our input regions resembles chromatin accessibility determined by DNaseI- or ATAC-seq at these sites (Fig. 1b, S1d). Accordingly, correlation analyses of genome-wide read distribution further confirmed a high correlation of our input library with DNaseI- and ATAC-seq profiles (Fig. 1c and S1e), validating that our library captured open regions, which are enriched for regulatory elements (Klemm, Shipony and Greenleaf, 2019), on a genome-wide scale.

**Figure 1:**
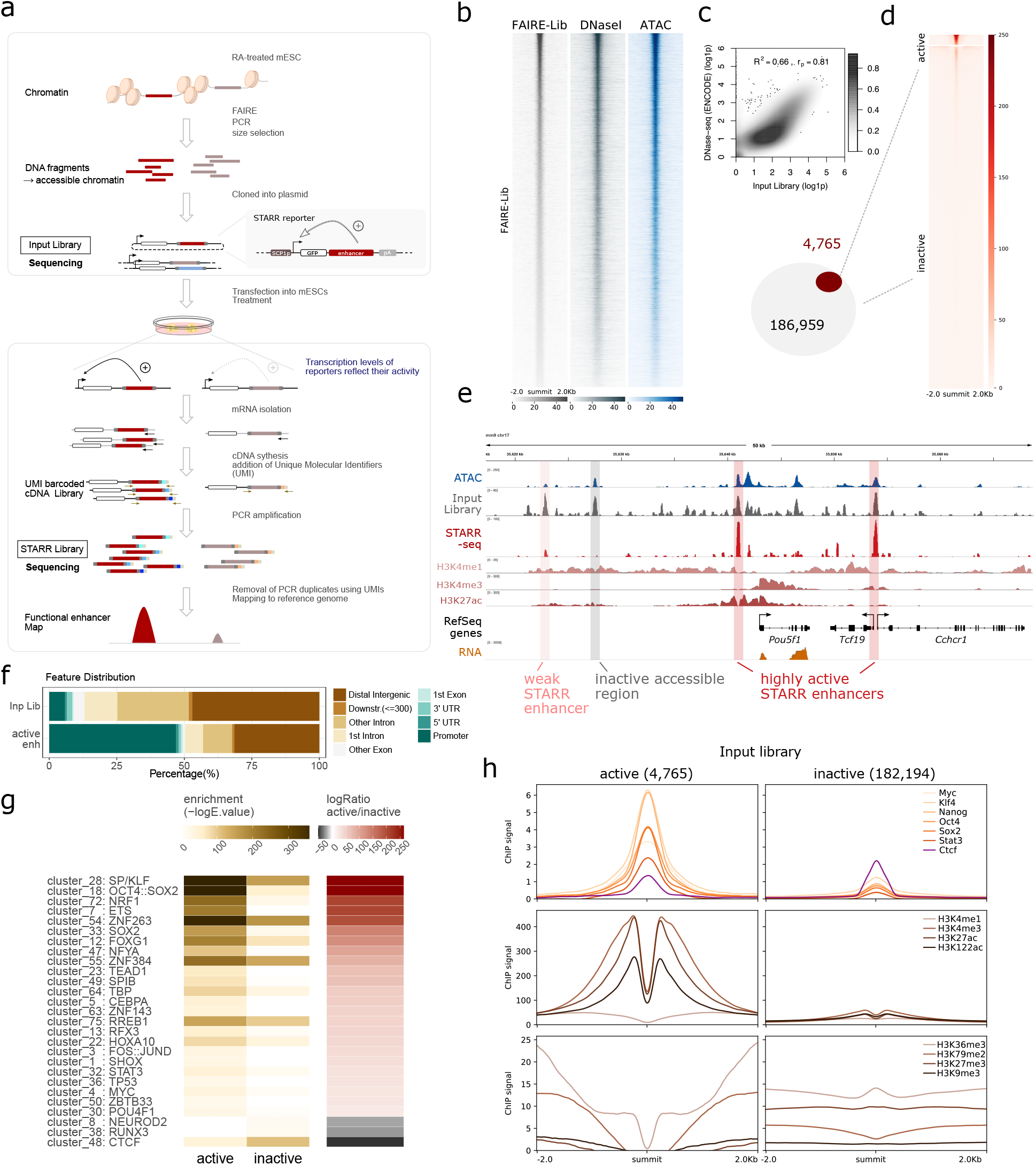
FAIRE-STARR-seq in mouse embryonic stem cells. a) Schematic representing the workflow of FAIRE-STARR-seq. b) Heatmaps depicting normalized read distribution of the FAIRE-STARR input library, DNase- and ATAC-seq at the FAIRE-STARR input regions. c) Correlation analysis of genome-wide read distribution, comparing the input library with DNase-seq data (ENCODE). Normalized and log1p transformed reads per 10 kb genomic bin are shown. d) Heatmap showing normalized FAIRE-STARR-seq signal at active (4,765) or inactive (182,194) input regions. e) Exemplary genomic region encompassing the *Pou5f1* gene. The FAIRE-STARR-seq signal merged from three replicates is shown and inactive, active, and highly active regions present in the library are highlighted. In addition, ChIP-seq data of histone modifications (HMs) as indicated, RNA-seq, ATAC-seq and input library signal from mESCs are shown. f) Genomic distribution of input regions and FAIRE-STARR enhancers with respect to annotated Refseq genes. Promoters were defined as the regions 1 kb upstream of a TSS. g) Motif enrichment analysis comparing the 4,765 FAIRE-STARR active and an equal number of randomly sampled inactive input regions. Enrichment of motif clusters is indicated as -log10E-value and the -log ratio comparing active versus inactive enrichment is shown. Enriched motifs (E <= 1e-5) with a minimum 20-fold -log difference of E-values between the two groups are shown. The JASPAR 2018 vertebrate clustered motif database was used as reference and listed TF names display TF groups clustered by consensus motif similarity (Khan *et al*., 2018). h) Anchorplots showing mean normalized ChIP-seq enrichment of the indicated HMs or TFs at FAIRE-STARR active or inactive input regions.

To get quantitative information regarding enhancer activity, we developed a modified version of the STARR-seq assay, which introduces unique molecular identifiers (UMIs) during the reverse transcription step (Fig. 1A). A similar approach has been described, however in that case UMIs are introduced in a first PCR cycle after the reverse transcription step (Neumayr *et al*., 2019). The introduction of UMIs allows one to distinguish between biological replicates and PCR duplicates that can dramatically distort the relative quantities of individual fragments within a library (Islam *et al*., 2014). Genome-wide correlation analyses of read distributions of individual FAIRE-STARR-seq samples revealed that outlier regions with extremely high read coverage in only one replicate were efficiently removed during the UMI filtering step (Fig. S1c) indicating that such regions are PCR amplification artefacts. Upregulation of interferon genes in response to transfection with plasmids can also distort STARR-seq reporter activation (Muerdter *et al*., 2018). To test if this is a potential problem in the mESCs used in our study, we analyzed the expression levels of selected interferon response associated genes. However, for each of the genes analyzed, the levels were below the qPCR detection limit regardless of whether the cells were transfected or not (data not shown) indicating that the interferon response is not activated upon transfection and thus should not influence the STARR activity read-out in our assays.

Next, active FAIRE-STARR enhancers were identified by performing peak calling for significantly enriched regions using the input library as background. This was done for each of the three biological replicates individually and for the merged replicates. We focused only on high-confidence regions by filtering for enhancers which were called in at least two replicates and were captured by at least three different fragments. Using these criteria, we identified 4,765 active enhancers with assigned quantitative STARR-scores, while the majority of the input regions showed no enhancer activity (Fig. 1d and e). To determine what distinguishes active enhancers from their inactive counterparts (182,194 accessible regions without STARR activity covered by our library), we analyzed sequence composition, TF occupancy and enrichment of a panel of histone modifications (HMs) linked to enhancers in mESCs (Creyghton *et al*., 2010; Dunham *et al*., 2012). In order to reduce the redundancy inherent in many motif databases that contain multiple highly similar motifs for related TFs, we used the JASPAR 2018 clustered vertebrate motif matrices which group related motifs into non-redundant clusters (Castro-Mondragon *et al*., 2017; Khan *et al*., 2018). As expected, we found that active enhancers are enriched for sequence motifs of pluripotency TFs such as POU5F1::SOX2 (cluster 18), SOX2 (cluster 33), MYC (cluster 4) and STAT3 (cluster 32) (Fig. 1g). Furthermore, CG-rich motif clusters (28: SP/KLF, 54: ZNF263, and 72: NRF1) were enriched for these regions. We also compared the quantity of enriched motifs between active and inactive regions and found that the number of significant motif hits is higher for active enhancers when compared to their inactive counterparts (Fig. 1g). On the other hand, inactive regions are characterized by an enrichment for motif clusters NEUROD2 (cluster 8) and RUNX3 (cluster 38), which contain TFs associated with differentiation and cell-type specific TFs, most of which are not expressed in mESCs (Fig. S1f) suggesting that these regions might be primed for activation when ESCs differentiate towards different cell types. Interestingly, the motif for CTCF, a master regulator of the genomic architecture (Phillips and Corces, 2009), is enriched for both groups, but this enrichment is more pronounced for inactive regions (Fig. 1g). Consistent with the observed motif enrichments, we found that ChIP-seq data for a panel of TFs involved in pluripotency showed higher levels of genomic occupancy at active enhancers than at their inactive counterparts whereas CTCF occupancy was slightly higher for inactive regions (Fig. 1h). To compare the chromatin landscape at the endogenous genomic loci between active enhancers and inactive regions, we performed ChIP-seq for eight HMs in mESCs. Intersection of the HM data showed that all three investigated HMs associated with active enhancers, H3K4me1, H3K27ac, and H3K122ac, as well as the promoter mark H3K4me3 are highly enriched at active-compared to inactive input regions. For HMs associated with transcription, H3K36me3 and H3K79me2, we also find an enrichment however not directly at the active enhancer but rather in the regions flanking it (Fig. S1h). In contrast, repressive marks H3K27me3 and H3K9me3 are depleted at active regions when compared to inactive input regions. Consistent with elevated H3K4me3 levels, we found that almost half of the active FAIRE-STARR enhancers are promoter-proximal regions. This percentage is much higher than in our library for which less than 10% of the regions map near promoters (Fig. 1f). These findings are consistent with a published study showing that promoters can act as enhancers that control the expression of other genes (Dao *et al*., 2017). Taken together, we established a quantitative approach to determine the enhancer activity of accessible genomic regions resulting in a genome-wide catalog of putative regulatory regions in mESCs.

### Quantitative FAIRE-STARR-seq identifies transcription factors associated with distinct enhancer activity levels

In addition to identifying which regions are active, the UMIs added during the reverse transcription step facilitate a quantitative assessment of enhancer activity based on the FAIRE-STARR data. This allowed us to rank the identified active FAIRE-STARR regions by their activity and, for example, to screen for features associated with different activity levels. To determine if enhancer activity correlates with expression of nearby genes, we first grouped the active regions into five consecutive quantiles of ascending activity (Fig. 2a). Next, for each group the individual regions were associated to neighboring genes by distance. Using this approach, we found that the expression levels for the genes of each category correlate with the enhancer activity scores with significant differences between the neighboring groups (Fig. 2b). These findings suggest that our quantitative FAIRE-STARR scores recapitulate the activity of enhancers in their endogenous genomic setting where they influence the expression level of nearby genes. Similarly, H3K27ac levels, a mark that is used as a proxy for enhancer activity (Shlyueva, Stampfel and Stark, 2014), correlate positively with our STARR score (Fig. S2a). This further indicates that our STARR activity scores capture the activity of enhancers in their endogenous genomic setting.

**Figure 2.**
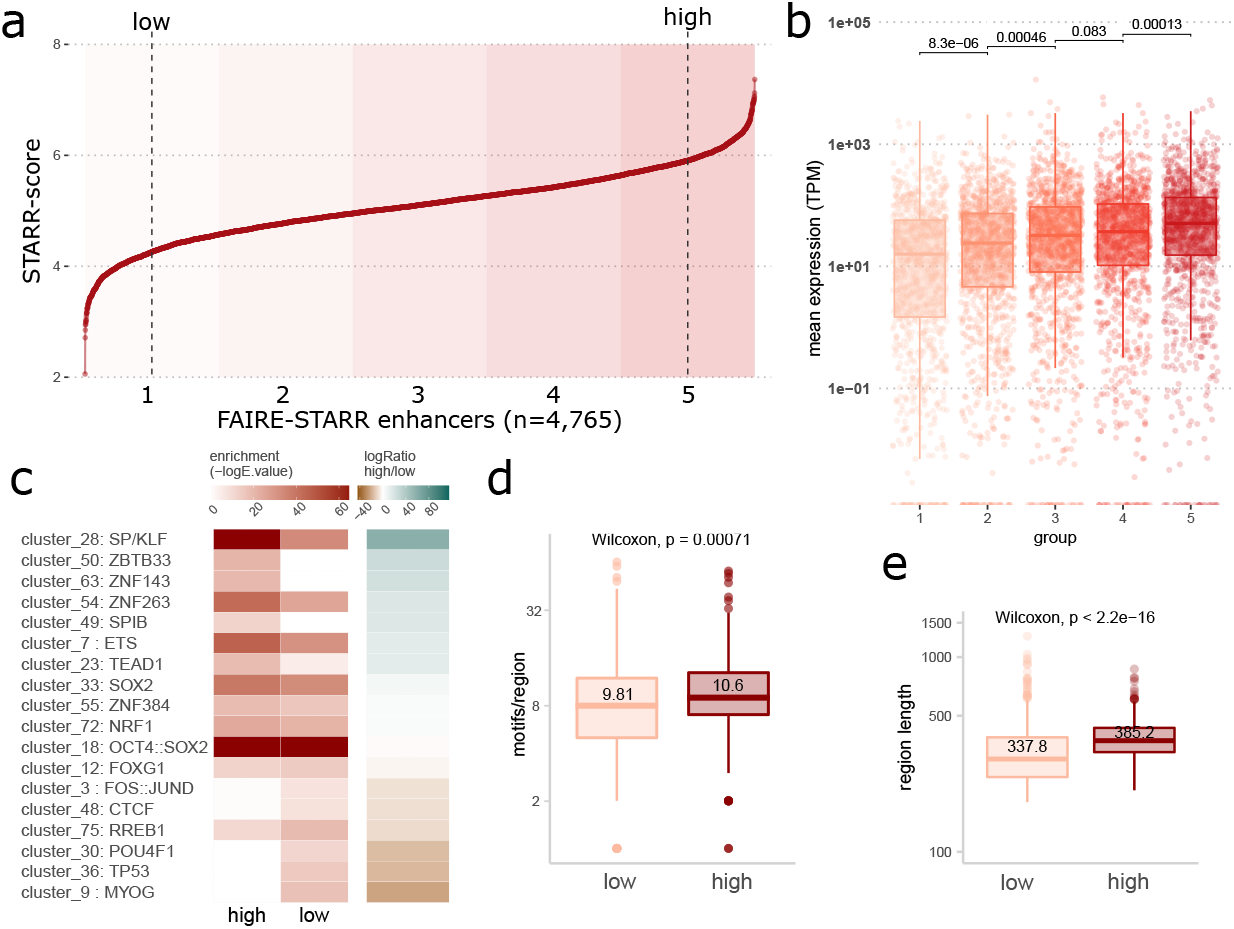
FAIRE-STARR-seq enables quantification of enhancer activity and activity level-associated sequence features. a) FAIRE-STARR enhancers were ranked for their activity (log STARR score) and divided into five groups of ascending enhancer activity (highlighted by increasing background coloring). Dashed lines depict the 10^th^ and 90^th^ percentiles of STARR activity. b) Expression of genes paired with FAIRE-STARR enhancers, for each of the five activity groups as depicted in a). Genes were paired with FAIRE-STARR enhancers by distance using GREAT (McLean *et al*., 2010) and TPM values of RNA-seq data are shown. Boxplots depict the distribution of expression of all genes per group, whiskers extend to 1.5 IQR. TPM values of individual genes are shown as dots. P-values for unpaired Wilcoxon tests comparing neighboring groups are indicated. c) TF sequence motifs enriched at active FAIRE-STARR enhancers, comparing the most active 10% (high) and least active 10% (low) of the active enhancers. Enriched motifs (E <= 1e-3) with a minimum 1-fold -log enrichment ratio between the two groups are shown. The JASPAR 2018 vertebrate clustered motif database was used as reference and a representative TF for each cluster is listed (Khan *et al*., 2018). Boxplots depicting d) the number of significantly enriched motifs and e) length of low- or high-ranking enhancers. Means are indicated as well as p-values for unpaired Wilcoxon tests comparing the two groups.

To investigate the role of DNA sequence in directing different levels of enhancer activity, we performed TF motif enrichment analysis comparing active enhancers that are ranked either at the top or bottom ten percent by STARR activity score (“high” and “low”, Fig. 2c). Interestingly, we found that the motifs for pluripotency TFs OCT4, SOX2 and NANOG (cluster 18) are enriched for high-as well as low-ranking enhancers indicating that high activity levels are not dictated by the presence of sequence motifs for these TFs. Rather, the top-ranking enhancers are characterized by motifs of the SP/KLF (cluster 28) and ETS (cluster 7) TF families. These factors are ubiquitously enriched at promoters, irrespective of the cell type and are accompanied by motifs for cell-type specific TFs (Landolin *et al*., 2010). For the low-activity enhancers, we found enrichment of motifs of cell type defining TFs, such as MYOG (cluster 9) and POUF4 TFs (cluster 30), suggesting a priming of enhancers that might play a role in later developmental stages when these TFs become expressed. Low activity enhancers were also enriched for the motif of p53, which was recently found to bind “dormant” enhancers in mESCs that are located in inaccessible chromatin and become activated upon cellular stress or during reprogramming (Peng *et al*., 2020). Another plausible explanation for increased activity levels of enhancers could be the absolute number of motifs per enhancer as well as on the diversity of these motifs (Singh *et al*., 2021). Accordingly, we found that the high-activity enhancers, on average, contain more motifs (10.6 compared to 9.8 average motifs/enhancer, Fig. 2d) and were 14% larger than enhancers with low activity (385 bp versus 338 bp for low-activity enhancers, Fig. 2e). Together, these findings indicate, that the nature of the sequence motifs present as well as the absolute number of motifs are critical drivers of enhancer activity.

### Epigenetic features and transcription factor occupancy define distinct enhancer subsets

Enhancers can be predicted based on the patterns of HMs present, with active enhancers harboring a high H3K4me1 to H3K4me3 ratio as well as high H3K27ac levels at flanking nucleosomes (Creyghton *et al*., 2010; Dunham *et al*., 2012; Ramisch *et al*., 2019). However, alternative histone marks for active enhancers have been described (Pradeepa *et al*., 2016; Martire *et al*., 2019; Armache *et al*., 2020). Moreover, although HMs correlate with enhancer activity it is largely unclear if this reflects a causative link (reviewed in Morgan and Shilatifard, 2020). To gain insight into the epigenetic landscape present at the active STARR enhancers, we clustered the active enhancers based on ChIP-seq signal for a panel of eight different HMs (Fig. 3a). Consistent with our finding that active enhancers are enriched in promoter regions (Fig. 1f), we found that about half of the active FAIRE-STARR enhancers show HMs characteristic for active promoters (cluster A: high H3K4me3, low H3K4me1, and high H3K27ac signal). An overlay with annotated promoter regions confirmed that the enhancers of cluster A (we called these “enhancer-promoters” or in short “E-promoters”) map to annotated promoters (Fig. 3a). Moreover, RNA-seq as well as H3K36me3 and H3K79me2 levels show that the genes at these promoters are actively transcribed in mECSs. Notably, enhancers of this cluster do not display the classical enhancer signature of high H3K4me1 over H3K4me3 levels, and consequently are not recognized by the CRUP enhancer prediction tool (Ramisch *et al*., 2019), which like many prediction tools prioritizes high H3K4me1 over H3K4me3 levels to call enhancers. This is different for enhancer clusters B to E which display high H3K4me1 levels and varying levels of H3K27ac, thus displaying a typical epigenetic signature of active enhancers and accordingly a larger overlap with enhancers predicted by CRUP. Clusters B and C, which display higher H3K27ac signals than clusters D and E, are highly active enhancers, showing high STARR activity as well as higher CRUP prediction scores. Cluster F, on the other hand, displays a typical H3K4 methylation pattern of enhancers but is lacking high H3K27ac levels, indicative of enhancers poised for activity (Creyghton *et al*., 2010; Rada-Iglesias *et al*., 2011). Accordingly, these enhancers have quite low enhancer prediction scores, but still can be identified as active in our FAIRE-STARR-seq assay. This indicates, that FAIRE-STARR-seq is able to pick up enhancers with a poised HM signature, while CRUP discards those regions by design.

**Figure 3.**
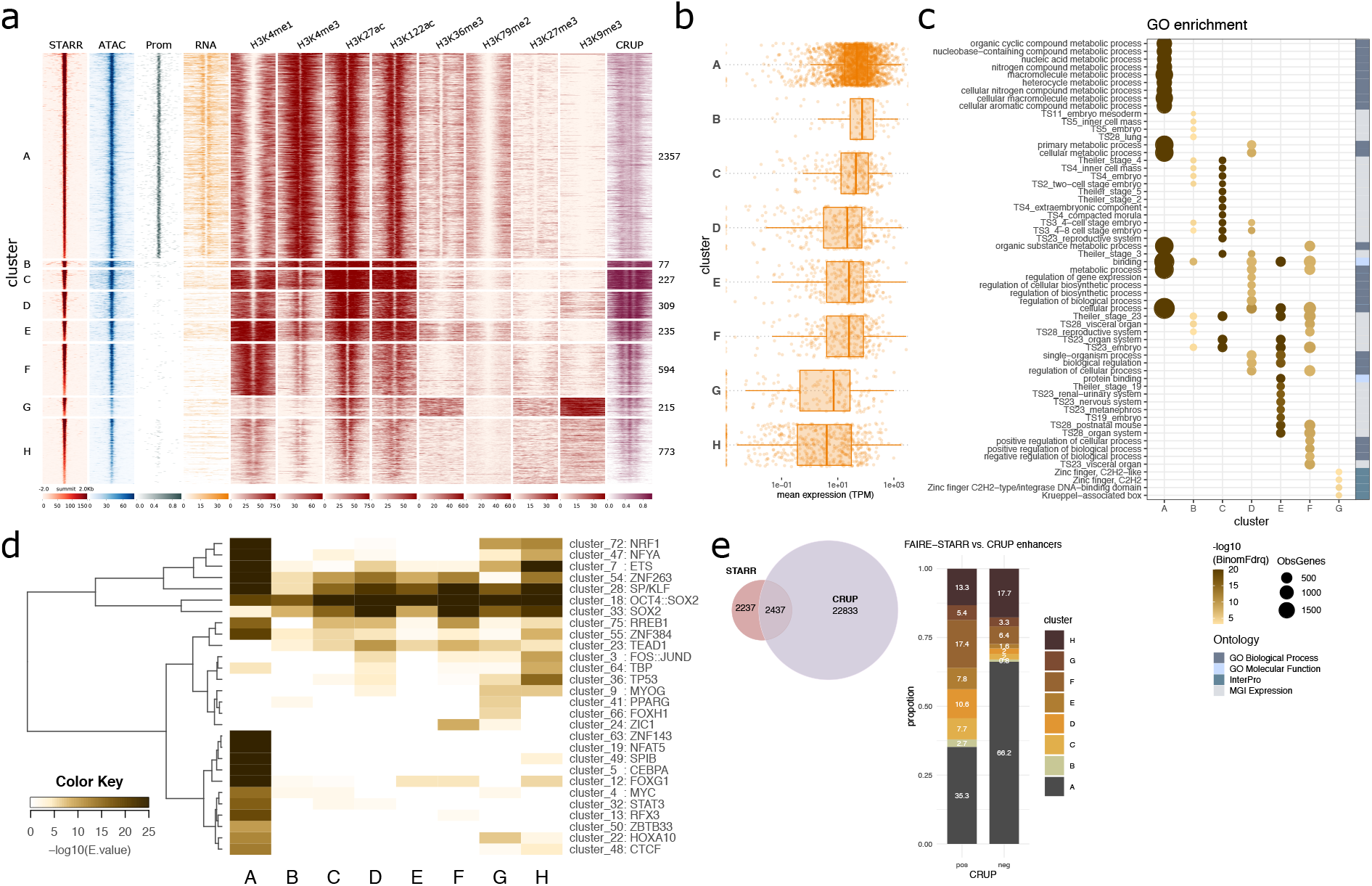
Functional mESC enhancers reside in different epigenomic environments. a) FAIRE-STARR enhancers were clustered (k-means clustering) based on the enrichment of the eight investigated histone modifications. For each cluster, the STARR-, ATAC-, and RNA-seq signals were plotted as were promoter regions (Prom) defined as one kb up- and downstream of the TSSs of annotated Refseq genes. Enhancer probability scores predicted by CRUP from mESC data are also shown. b) Genes were assigned to enhancer clusters using GREAT and RNA-seq expression data is shown as dots for individual genes (TPM normalized) and as boxplots for each enhancer cluster. c) Gene ontology analysis of genes associated with each enhancer cluster showing the fifteen most significant GO terms per cluster and their false discovery rate (-log10BionomFdrQ, cutoff 1e-03). For each ontology, the number of observed genes (ObsGenes), the significance, and the source of the assigned ontology are shown. d) TF motif enrichment analysis (AME) for each enhancer cluster using the JASPAR 2018 vertebrate clustered motif matrices. TF motifs which were enriched (E <= 1e-5) for at least one cluster were clustered for TF occurrences applying wards clustering and Manhattan similarity measures. e) Venn diagram showing the intersection of FAIRE-STARR and CRUP enhancers. FAIRE-STARR enhancers which overlap with CRUP enhancers (pos) or do not overlap (neg) were assigned to the HM clusters defined in a.

Cluster G, shows a rather uncommon HM pattern of enriched H3K36me3, a mark for active transcription, together with elevated H3K9me3, a hallmark of heterochromatin. The combination of these two marks has previously been reported to occur at the same nucleosome to mark lowly expressed genes and weak enhancers (Mauser *et al*., 2017). Interestingly, alignment with the RepeatMasker database (Smit AFA, Hubley R, 2013) revealed that 93% of cluster G enhancers map to repeat elements. The majority of these repeats are from the endogenous retrovirus-K (ERVK) family (Fig. S3f), a rather young family of mouse-specific endogenous retroviruses, which can act as enhancers in mESCs (Sundaram *et al*., 2017). Finally, cluster H, which exhibits the lowest STARR signal, also shows the lowest accessibility based on our ATAC-seq data and lowest enrichment of each HM except H3K9me3. This indicates that these regions are not very accessible, nor active, in the endogenous genomic setting and may only be able to unleash their activity in the episomal STARR-seq setting where such sequences are taken out of their repressive endogenous chromatin context.

Motivated by a study claiming that H3K122ac marks a unique class of active enhancers lacking H3K27ac (Pradeepa *et al*., 2016), we included this mark in our ChIP-seq experiments. However, contrary to the published study, we did not observe a convincing cluster with H3K122ac but lacking H3K27ac. Rather, we found that, in general, the signal for H3K27ac and H3K122ac is essentially the same at enhancers. This is different for promoters, where we found H3K122ac enriched at H3K4me3-marked gene promoters, irrespective of the H3K27ac state (Fig. S3h).

To study the link between enhancer clusters and nearby gene expression and to test if they are associated with different categories of genes, we paired the clustered enhancers with genes by distance and analyzed gene expression levels (Fig. 3b) as well as gene ontologies (Fig. 3c). Overall, we found that the expression of enhancer cluster-associated genes correlates well with epigenetic signatures of individual clusters. For example, we find the highest mean expression level for genes associated with clusters B and C, that show the most prominent signatures of active enhancers (Fig. 3b). In contrast, we find the lowest expression levels for genes associated with clusters G and H, two enhancer clusters with low levels of active enhancer marks. Interestingly, the function of the genes associated with different clusters also diverges. For instance, E-promoter cluster A-associated genes are involved in more general cellular processes, such as nucleic acid, nitrogen compound, and organic substance metabolic processes, whereas genes associated with clusters B and C play a role in early embryonic stages (Fig. 3c). Enhancer clusters D to F are associated with genes of medium expression levels, which are enriched for genes typically expressed at later time points during embryonal development. Cluster G enhancers correspond to genes with rather low expression levels which are enriched for zinc finger and KRAB-domain containing genes. The genes of this family of TFs have been described as marked by H3K36me3 and H3K9me3 (Valle-García *et al*., 2016), the combination we now identify at the cluster of associated enhancers as well. Finally, enhancer cluster H is associated with genes with the lowest mean expression levels of all investigated clusters and no significantly enriched GO terms could be identified. This is in line with our hypothesis that these enhancers are a heterogenous group, which are repressed in mESCs but can be activated at different points during differentiation.

Next, we compared the sequence composition of the enhancer clusters, and found that they display characteristic patterns of motif enrichment (Fig. 3d). One example is the E-promoter cluster A, which is enriched for many motifs including CG-rich motifs like SP/KLF family (cluster 28) and NRF1 (cluster 72), as well as motifs of the ETS family (cluster 7). Similarly, analysis of published TF ChIP-seq data showed different binding patterns for individual enhancer clusters (Fig. S3a). Consistent with a selective enrichment of their motifs at E-promoters, we found that c-MYC and Ronin selectively occupy enhancers of E-promoter cluster A (Fig. S3a). The situation is different for KLF4 (a member of the SP/KLF family) which, as expected, binds E-promoter cluster A, but also to other enhancer clusters that are enriched for the SP/KLF motif (Fig. S3a). The motif for pluripotency factors OCT4 and SOX2 (cluster 18) is enriched for all enhancer clusters, with the lowest enrichment for cluster A, and ChIP-seq showed preferential binding at clusters B to F. Enhancer cluster F, lacking the active mark H3K27ac, is enriched for TF motifs of the ZIC family of TFs (cluster 24) that are implicated in the transition from naïve to primed pluripotency (Yang *et al*., 2019). Finally, p53, TBP and FOS:JUN motif clusters (36, 64, and 3) were specifically enriched for repressed enhancer cluster H and p53 was recently described to bind and activate repressed enhancers upon stress or differentiation stimulus (Peng *et al*., 2020).

Of note, apart from the transcriptionally active E-promoters, we also find many actively transcribed promoters without FAIRE-STARR-seq activity (Fig. S3b) showing that enhancer activity is not a general feature associated with active promoters. Comparison of these promoters with E-promoters shows stronger enrichment of c-MYC and RONIN at E-promoters, but similar KLF4 occupancy at both groups (Fig. S3c and d). Further, motif enrichment analysis comparing E-promoters and promoters lacking STARR-seq activity revealed differences in sequence composition (Fig. S3e). For example, consistent with the observed selective enrichment by ChIP, the motif for MYC (cluster 4) was more highly enriched for E-promoters than for regular promoters, while motifs for pluripotency factors OCT4, SOX2, and NANOG (cluster 18) were more enriched at regular promoters that at E-promoters.

A global comparison between enhancers predicted based on chromatin features using CRUP and active enhancers identified by FAIRE-STARR revealed that only 2,437 (52.1%) of the active STARR enhancers were also predicted by CRUP (Fig. 3e) while CRUP predicted 22,833 regions that were not identified by FAIRE-STARR. The majority of FAIRE-STARR enhancers which were not predicted by CRUP fall into cluster A, the E-promoters, and thus display a chromatin signature which is filtered-out by CRUP. The second largest group of STARR enhancers not detected by CRUP are cluster H regions, repressed enhancers, which are not marked by the classical enhancer signature recognized by CRUP. On the other hand, enhancers which were only predicted by CRUP but not picked-up by FAIRE-STARR-seq showed very low chromatin accessibility by ATAC, and overall lower activation marks (Fig. S3g). This indicates, that these regions were not included in the FAIRE-STARR input library and thus could not be identified as active.

Together, we find that enhancers can be grouped based on different HM patterns. These enhancer clusters have different activities and are associated with genes with different functions. The partial overlap with CRUP enhancer predictions highlights that HM-based enhancer predictions and functional assays such as FAIRE-STARR-seq can provide complementary information. For example, FAIRE-STARR-seq can be used to identify enhancers with typical enhancer signatures, but also to find E-promoters, poised, and repressed enhancers, which exhibit activity in the episomal reporter background but are not picked-up by enhancer predictions. Additionally, our data is in line with other studies suggesting that a formal distinction between promoters and enhancers, and the exclusion of promoter signatures from HM-based predictions, might not make sense given that promoters can exert both functions (Kim and Shiekhattar, 2015; Dao *et al*., 2017; Dao and Spicuglia, 2018).

### Sequence-based prediction of active enhancers in mESC

As shown above, enhancer prediction based on HMs often excludes E-promoters and depends on several ChIP-seq data sets which are not always available. Here, we set out to build an enhancer prediction model solely based on DNA sequence using their activity scores from FAIRE-STARR-seq. Given the different sequence composition and accordingly motif enrichment of promoter and enhancer regions (Fig. 3d), we chose to analyze candidate regions that map to these two regions separately. Specifically, we took all accessible regions from our input library and divided them into two groups: Those overlapping with annotated promoters and those that do not overlap. Next, within each group, we ranked the regions by their STARR-score and used the 1% or 10% of regions with the highest or the lowest STARR-score to train two regularized logistic regression (elastic net) models to classify active and inactive DNA sequences (Fig. 4a, b, and c). As features for each model, we used the width of the region and enrichment of clustered JASPAR motif matrices (Castro-Mondragon *et al*., 2017). Each elastic net regression classifier was fitted to maximize the cross-validated mean AUC which yielded 0.75 for enhancer regions (Fig. 4b) and 0.87 for E-promoters (Fig. S4a). Thus, the classifier for both enhancers and E-promoters performed quite well, suggesting that enrichment patterns of TF motifs alone can be used to distinguish between active and inactive regions for both promoter and enhancer regions with reasonable accuracy.

**Figure 4:**
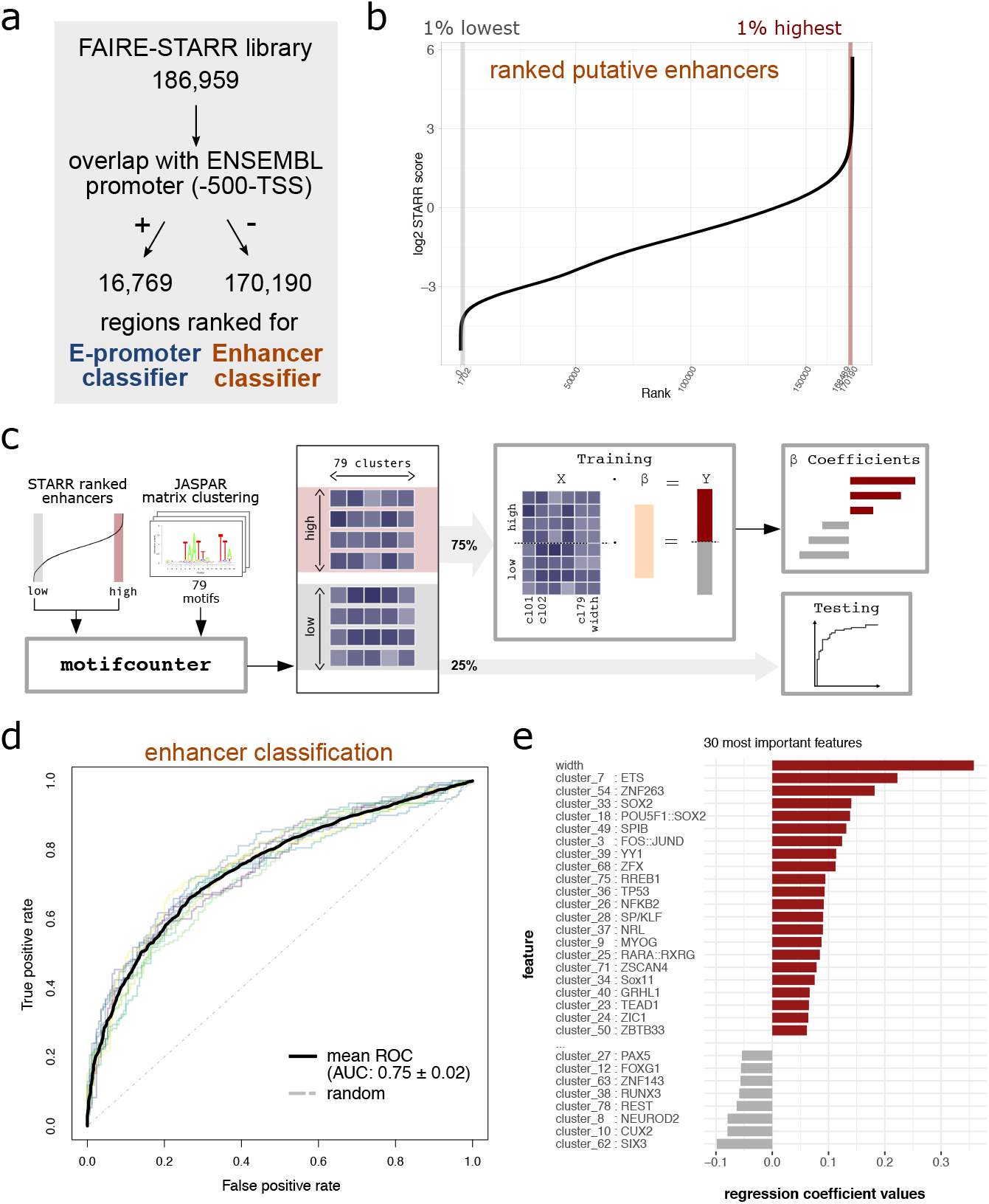
Sequence-based prediction of active enhancers. a) The regions of the FAIRE-STARR library were first divided by their overlap with ENSEMBL promoters (region up to 500 bp upstream of a TSS). Regions which overlap with promoters were used to generate an E-promoter prediction model, whereas those not overlapping were used for the enhancer prediction model. To this end both groups (b) putative enhancers and S4a) promoters) were ranked for their STARR score, and 1 or 10% highest or lowest ranking regions were used for model generation. c) Cartoon depicting how the enhancer prediction model was trained on ranked regions from our FAIRE-STARR-seq analysis using enrichment of JASPAR 2018 vertebrate clustered motif matrices and region width as independent variables. 75% of the highest and lowest ranking regions were used for model training, while the remaining 25% were used for testing. d) Plot shows model performance as receiver operating characteristic (ROC) curve for each of the outer cross-validation folds, mean ROC curve with area under the curve (AUC) and its standard deviation. e) The 30 most predictive variables for the optimal enhancer prediction model and their coefficients are shown. Positive coefficients indicate a positive association with high STARR scores, while motifs with negative coefficients are associated with low-scoring elements.

To determine which features were most important in predicting if a region is active, we analyzed the ranked model coefficients (Fig. 4e and S4c) which reflect the importance of individual features for each optimized activity-prediction model. Interestingly, the two features with the highest coefficient for both the E-promoter and the enhancer-prediction model are enhancer width and the motif for ETS TFs (cluster 7). For classification of enhancers, the motif for pluripotency TFs SOX2, OCT4 and NANOG (cluster 18) was among the top features associated with active regions (Fig. 4e), while it showed a negative coefficient in the E-promoter classification (Fig. S4c), indicating that pluripotency factors contribute to enhancer but not E-promoter activity. Similarly, the motif for CTCF scored a positive coefficient for the E-promoter classification (Fig. S4c), while it was slightly negative for active enhancers (not in figure, coeff. = -0.045).

Together, our modeling demonstrates that for both enhancers and E-promoters activity can be predicted from DNA sequence using a rather small feature set of 79 clustered TF motifs and enhancer width as input.

### FAIRE-STARR-seq identifies enhancers that change their activity upon exiting pluripotency

During ESC differentiation, pluripotency genes are gradually shut down, while genes important for early differentiation and later cell-type specific genes become activated (Young, 2011; Semrau *et al*., 2017). This process is accompanied by gain and loss of activity of differentiation- and pluripotency-specific enhancers, respectively. To identify enhancers that change their activity upon exiting pluripotency, we analyzed enhancer activity during the early stages of differentiation using the FAIRE-STARR-seq approach. Specifically, we compared transfected cells treated with LIF to maintain pluripotency with cells from the same transfection-batch that were treated for 4 hours with retinoic acid (RA) to initiate differentiation towards the neuronal lineage (Fig. 5a) (Gudas and Wagner, 2011). We then focused on enhancer regions which either lost (LIF-dependent) or gained (RA-inducible) activity upon differentiation. This resulted in the identification of 616 LIF-dependent and 386 RA-inducible enhancers, which show varying degrees of loss or gain of STARR activity (Fig. 5b). The activity of these enhancers correlated with changes in the expression of nearby genes, with genes near RA-inducible enhancers showing, on average, an increase in gene expression (and more associated upregulated than down-regulated genes) whereas the expression genes near LIF-dependent enhancers showed, on average, a slight decrease in expression upon differentiation (and more repressed than upregulated genes, Fig. 5c and S5b, c). Notably, when compared to all active mESC enhancers (Fig. 1f) that are typically promoter-proximal, the differentially active enhancers are most frequently found at distal intergenic regions (Fig. S5a) and display distinct TF motif enrichments (Fig. 5d). For instance, RA-inducible enhancers are more enriched for RARα::RXRα and ETS-family motifs than the LIF-dependent enhancers consistent with the activation of RAR upon treatment of cells with its cognate ligand. Accordingly, when we performed ChIP-seq targeting RARα from RA-treated cells, we found that RARα is enriched at RA-inducible enhancers whereas no such enrichment was found for the LIF-dependent enhancer regions (Fig. S5d). Similarly, consistent with STAT3 activation by LIF, STAT3 binding is more enriched at the LIF-inducible enhancers than at RA-inducible enhancers (Fig. S5d). Moreover, LIF-dependent enhancers are more enriched for OCT4:SOX2, SP1-like family, SOX2, NFY, TEAD and NRF1 motifs than the RA-inducible enhancers. The enrichment of these sequence motifs is reflected in enriched binding of OCT4, SOX2, NANOG, and KLF4 based on published ChIP-seq data (Chen *et al*., 2008, Fig. S5d). Interestingly, binding of OCT4, SOX2, and NANOG was not only enriched at LIF-dependent but also at RA-inducible enhancers when compared to all active mESC enhancers. This indicates that pluripotency TFs play a supportive role at RA-inducible enhancers during the early stages of differentiation.

**Figure 5:**
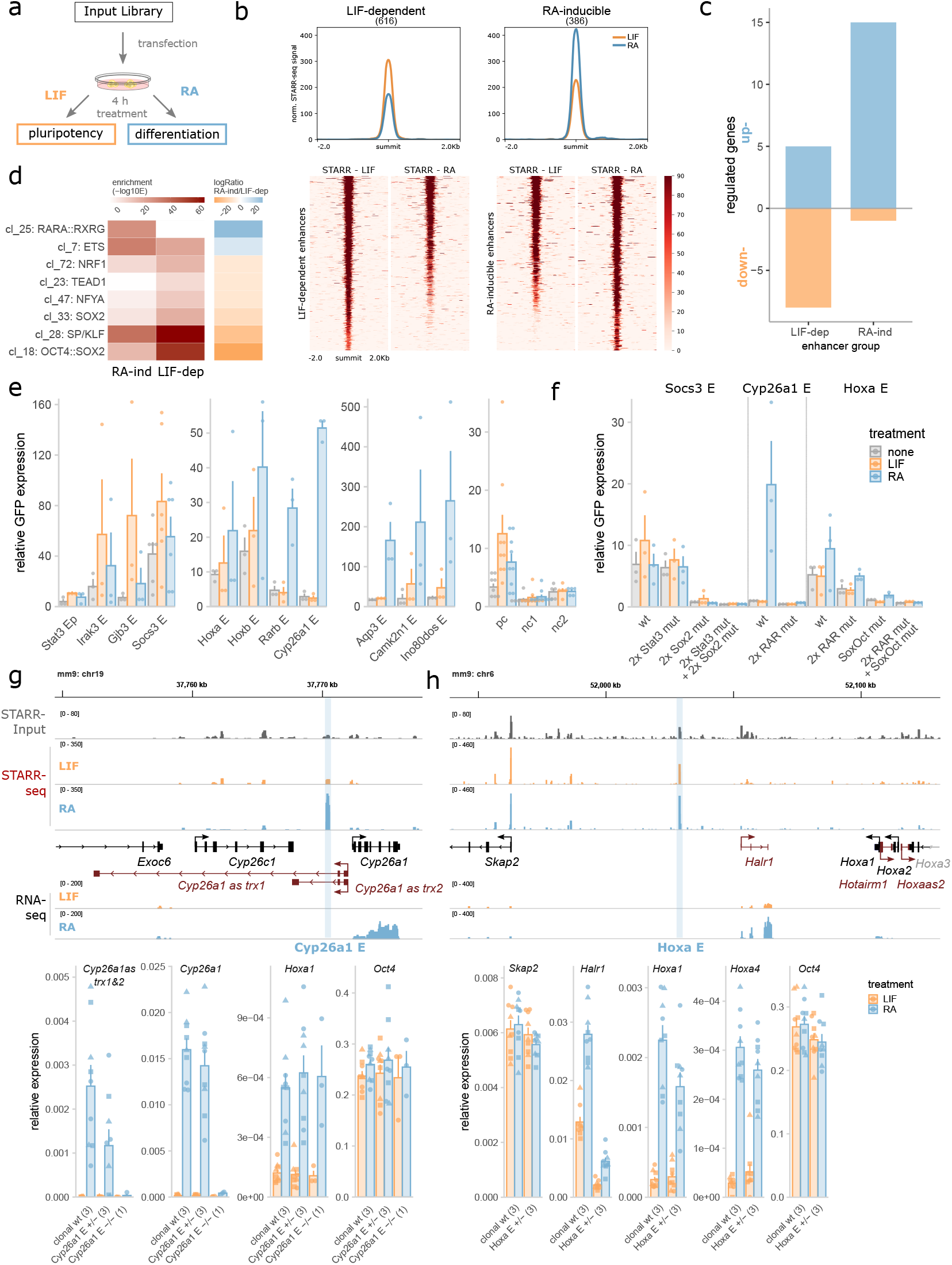
Differentiation-associated changes in enhancer activity. a) Treatment scheme to investigate how inducing differentiation of mESCs changes enhancer activity. b) Mean FAIRE-STARR signal (top) and heat-maps (bottom) for LIF-dependent and RA-inducible STARR enhancers. c) Number of differentially expressed genes (|log2FC| >= 1, p.adj <= 0.05) paired with enhancers by distance using GREAT. d) Differentially enriched TF motif clusters (JASPAR 2018 vertebrate clustered matrices) for RA-induced and LIF-dependent enhancers were identified using AME (E <= 1e-3, -logRatio >= 5). e) Candidate FAIRE-STARR enhancers were cloned individually and assessed for enhancer activity by RT-qPCR targeting GFP reporter transcripts. 20 h after transfection, E14 mESCs were treated for 4 h either with LIF, RA, or ES medium only (none). Bar plots show normalized mean expression ± SE of three replicates (dots). f) TF motifs-matches identified by JASPAR scan (Table S1) for enhancers as indicated were deleted by site-directed mutagenesis and activity was analyzed as described in e. g) and h) upper panels show genomic loci encompassing STARR enhancers selected for genomic deletion using CRISPR/Cas9. Normalized FAIRE-STARR-input, -seq, and RNA-seq signals are shown. RefSeq genes are shown in either black (protein coding genes) or red (non-coding genes). Lower panels depict the RNA expression of genes near the deleted enhancer and of control genes for clonal wild type (wt) and enhancer heterozygous (E +/-) and homozygous (E -/-) deletion clones. Bar plots represent mean gene expression ± SE of three biological replicates (dots) and 1-3 clonal cell lines (number indicated in brackets) after 4 h of LIF or RA treatment. Data points for individual clonal lines are shown as dots with matching shapes.

Next, the regulatory behavior of several LIF-dependent and RA-inducible enhancers identified in our screen was tested by cloning individual active regions into the STARR-seq vector and analyzing their activity by qPCR (Fig. 5e). One exemplary LIF-dependent enhancer we tested is located distal to the *Socs3* (suppressor of cytokine signaling-3) gene, which is activated by LIF-mediated STAT3 signaling (Yu *et al*., 2017). We confirmed that the *Socs3* enhancer is LIF-inducible and this induction is blunted when the two identified STAT3 motifs were mutated (Fig. 5f). Furthermore, mutation of the two SOX2 motifs of the *Socs3* enhancer leads to a marked loss of basal activity. Finally, the combined mutation of all SOX2 and STAT3 motifs abolished enhancer activity completely. This finding illustrates the importance of these TFs in facilitating LIF-dependent activity as suggested by the enrichment of sequence motifs for these TFs at LIF-dependent enhancers (Fig. 5d). We also characterized an RA-inducible enhancer upstream of the RA target gene *Cyp26a1*, which is pivotal for proper differentiation (Abu-Abed *et al*., 2001). The *Cyp26a1* enhancer is inactive during pluripotency but is massively upregulated upon differentiation (Fig. 5e, f). Consistent with a role of RAR in activating this enhancer, mutation of both of the identified RARα::RXRα motifs resulted in a complete loss of induction upon RA treatment (Fig. 5f). As a final enhancer, we analyzed the RA-inducible *Hoxa* enhancer (Fig. 5e, f), which is located over 70 kb upstream of the RA-responsive *Hoxa* gene cluster (De Kumar *et al*., 2015). The *Hoxa* enhancer shows basal activity in pluripotency which increases upon differentiation (Fig. 5f). Interestingly, mutation of the two identified RARα::RXRα motifs reduced basal activity during pluripotency but did not impair the RA-induced activation, suggesting that other motifs mediate the activation. As expected, mutation of the only identified OCT4:SOX2 motif of the *Hoxa* enhancer reduced activity in pluripotency whereas the combined mutation of both RARα::RXRα and the OCT4:SOX2 motifs completely abolished basal as well as RA-induced activation of the *Hoxa* enhancer. To test the role of two RA-inducible enhancers in the regulation of nearby genes in the endogenous genomic context, we generated CRISPR/Cas9-mediated genomic deletions of these enhancers in mESCs. The first enhancer we targeted was the *Cyp26a1* enhancer (E) described above (Fig. 5g). We were able to generate a single homo- and three heterozygous clonal lines for the *Cyp26a1* E deletion (Fig. S5e). Analysis of the clonal lines revealed that heterozygous deletion of the *Cyp26a1* E did not lead to an apparent impairment in upregulation of *Cyp26a1* whereas the homozygous enhancer knock-out led to a complete loss of inducibility (Fig. 5g). Interestingly, our RNA-seq data showed RA-inducible expression of two unannotated, spliced, and poly-adenylated transcripts, located anti-sense and only a few hundred basepairs upstream of *Cyp26a1* (*Cyp26a1 as trx1&2*). For these enhancer-proximal transcripts we found that RA-induction was impaired in the heterozygous lines whereas activation was completely lost in the homozygous *Cyp26a1* E deletion clone. Inducibility of *Hoxa1* by RA and expression of OCT4 (*Pou5f1*), two genes located on other chromosomes than *Cyp26a1* E, was not affected by the *Cyp26a1* E deletion, indicating that RA-signaling is still functional in the homozygous enhancer knockout and that the effects observed are specific for transcripts that are enhancer-proximal. For the other investigated enhancer, upstream of the *Hoxa* gene cluster (*Hoxa* E), we were able to generate only heterozygous deletion mutants (Fig. S5e). These mutants were still able to activate *Hoxa1* and *Hoxa4* upon RA treatment to induce differentiation, however with slightly reduced levels compared to wildtype indicating that this enhancer might contribute to the RA-dependent upregulation of the *Hoxa* gene cluster (Fig. 5h). The impact of the *Hoxa* E deletion was more prominent for the non-coding RNA *Halr1*, which is located upstream of the *Hoxa* genes and thus closer to the investigated enhancer. Specifically, the heterozygous deletion of the *Hoxa* E resulted in a marked decrease in *Halr1* expression during pluripotency and much lower levels when cells were treated with RA to stimulate differentiation. In contrast, expression of *Skap2*, a gene upstream of the *Hoxa* gene cluster, and mESC marker *Pou5f1* are not impacted by the *Hoxa* enhancer deletion, which indicates specificity of the observed effects. Thus, consistent with the data for the episomal reporter (Fig. 5f), this indicates a dual function of this enhancer to facilitate basal *Halr1* expression during pluripotency as well as induced expression upon differentiation.

Altogether, these results show, that the FAIRE-STARR-seq assay can be used to trace the dynamics of enhancer activity and can be used to identify enhancers which gain, but also those that loose enhancer activity upon induced differentiation. These enhancer subsets are characterized by distinct motif enrichments and the binding of specific TFs and are associated with regulation of nearby genes.

### Sequence features associated with RARα-occupied enhancers that change activity upon ligand binding

Our ChIP-seq experiments targeting RARα, the receptor of RA and key effector in RA-induced differentiation, uncovered thousands of binding sites. However, only a subset of these RARα-occupied sites show a change in activity upon RA-treatment in our STARR-seq experiments (Fig. 6a). Moreover, depending on the RARα-occupied region examined, we found that enhancer activity can either stay the same, go up, or go down upon addition of RAR’s cognate ligand RA (Fig. 6a, b). To identify sequence features that may play a role in determining if an RARα-occupied site changes its activity upon RA treatment, we first determined the effect of RA treatment on enhancer activity for all RARα-occupied sites covered in our STARR-seq library (Fig. S6a). Next, we defined three categories: Active RARα-occupied enhancers which (1) do not change activity upon RA treatment (“non-responding)”, (2) become more active upon RA treatment (“induced”) and finally, (3) RAR-occupied enhancers whose activity decreases upon treatment (“repressed”, Fig. 6b). Comparison of the motif composition of these three categories of RARα-occupied enhancers showed several differences that could play a role in determining if RA treatment induces a change in enhancer activity. For example, RARα::RXRγ heterodimer motifs (cluster 25) are more enriched for RARα-occupied regions that are activated in response to RA than either regions that are not regulated or those with repressed enhancer activity (Fig. 6c). Furthermore, clustered TF motifs of nuclear receptors, such as RXRA::VDR (cluster 2) and PPARG (cluster 41), and motifs for pluripotency factors (OCT4, SOX2, and NANOG, cluster 18) and SP/KLF (cluster 28) were more significantly enriched for induced RARα binding sites. On the other hand, RARα-occupied sites that lose activity upon RA treatment had the lowest enrichment of the canonical RARα::RXRγ heterodimer motif and were characterized by a relatively high occurrence of motif clusters ZNF384 (cluster 55), HOXA10 (cluster 22), and CTCF (cluster 48), which could thus play a role in RARα-dependent repression of enhancer activity. Side-by-side comparison of inducible, non-responding, and repressed RARα sites showed that RARα occupancy was comparable for inducible and non-responding regions with only slightly lower occupancy at repressed sites (Fig. 6f). Interestingly, chromatin accessibility assessed by ATAC was comparable for induced and repressed RARα-occupied sites but higher for non-responding sites in pluripotency and only increased marginally at induced sites upon RA treatment (Fig. 6f). Similarly, H3K27ac enrichment in pluripotency was lower at inducible and repressed than at non-responding RARα sites and inducible regions only reached comparable levels to repressed sites upon RA treatment (Fig. 6f). Accordingly, the enrichment of RARα and H3K27ac at RARα sites did not correlate positively with RA-inducibility (Fig. S6e), indicating that enhancer inducibility of an RARα site cannot simply be inferred from ChIP enrichment. This further indicates that sequence composition acts as an additional regulatory layer to control not only if an enhancer changes its activity but also the direction in which enhancer activity is modulated upon RA treatment.

**Figure 6:**
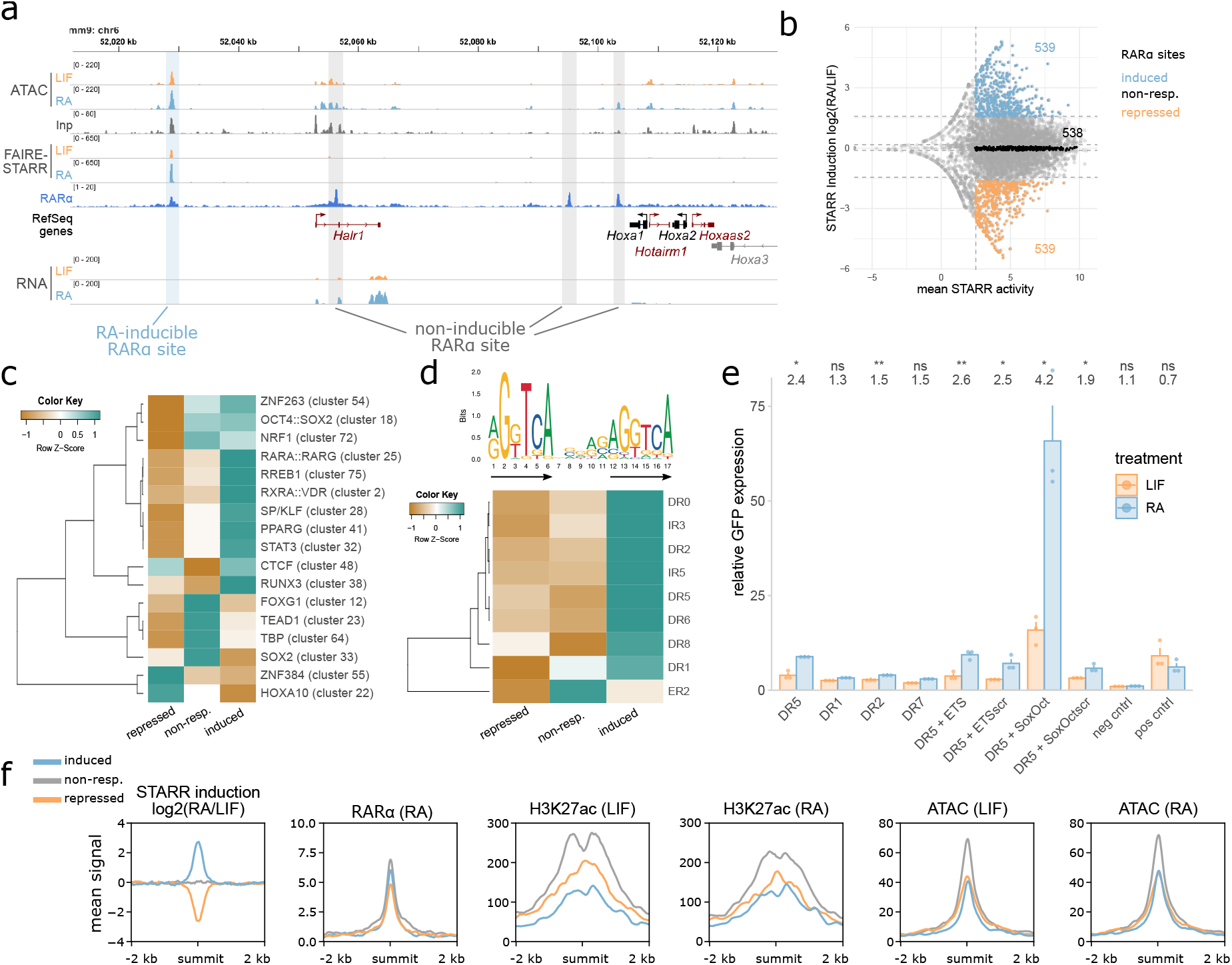
RA-induced changes in enhancer activity at RARα-occupied sites correlate with specific sequence and chromatin features. a) Genome browser view of an exemplary genomic region encompassing RA-inducible and non-inducible RARα binding sites. Normalized ATAC-, FAIRE-STARR and RNA-seq signals for LIF or RA treated cells and RefSeq genes for this region in either black (protein coding genes) or red (non-coding genes) b) Distribution of changes in STARR activity (log2 fold change STARR score RA/LIF) and mean STARR activity (log score, for both treatments) of RARα-occupied regions that are covered in our FAIRE-STARR input library (as shown in Fig. S6a). Only regions with a minimum mean STARR activity >= 2.5 were included for further analysis. The 10% most induced, 10% most repressed and an equal number of regions that do not respond to RA treatment (non-resp.) were used for motif enrichment and TF binding analyses. c) Enriched TF motif clusters (JASPAR 2018 clustered motif matrices) at induced, repressed, and non-responsive RARα-occupied sites. TF motif clusters with a maximum E-value of 1e-5 for at least one group and a log fold change >= 2 of induced or repressed over non-responsive regions are shown. Z-score normalization of E-values per row was performed. d) Different spacings (0-8 nucleotides) and orientations (direct (DR), inverted (IR), and everted repeat (ER)) of the RARα::RXRα consensus motif (MA0159.1, upper panel, arrows highlight repeat orientation) were constructed *in silico* and used for motif enrichment analysis using AME. Only motifs which showed significant enrichment (E-value <= 1e-3) for at least one RARα binding site group are shown. Z-score normalization of E-values per row was performed. e) Enhancer activity measured by STARR-RT-qPCR for selected spacing variants of the RARα::RXRα consensus motif (MA0159.1) and neighboring TF motifs as indicated (scr = scrambled motif) after 4 h of LIF or RA treatment. Bar plots depict the mean GFP expression + SE for three biological replicates. f) Mean enrichment of RARα and H3K27ac as well as chromatin accessibility (ATAC) at induced, repressed, and non-responsive RARα-occupied sites.

RARα typically binds as a heterodimer together with retinoic x receptor (RXR) to retinoic-acid response elements (RAREs) that can have distinct motif architectures, depending on cellular background and differentiation stage (Delacroix *et al*., 2010; Chatagnon *et al*., 2015). These motifs share the same consensus hexameric direct repeat (DR) however they differ in terms of orientation and the spacing between the repeat elements (Moutier *et al*., 2012). To elucidate the possible role of different spacings of the RARα::RXRα consensus motif (MA0159.1) on RA-induced changes in enhancer activity, we constructed different repeat orientations and spacings *in silico* (DR, everted (ER) and inverted repeats (IR) of the consensus motif with spacing from 0-8 nucleotides) and compared inducible, non-responsive, and repressed RARα-occupied regions for enrichment of these motifs (Fig. 6d). As previously described (Moutier *et al*., 2012), we find DR0 to be the most enriched spacing for RARα-occupied sites for all three enhancer subgroups (data not shown). Moreover, induced RARα-occupied regions are more enriched for each of the investigated motif architectures than their repressed counterparts and display higher abundance of the consensus repeat half-site (Fig. S6b) indicating that activation might be driven by direct RARα binding to its response element whereas repression is not. To determine how DR spacing influences RA-dependent regulation of enhancer activity in mESCs, we cloned single DRs with different spacings, but the same DR sequence into the STARR vector. Consistent with previous studies (Moutier *et al*., 2012), we found that activation was most prominent for the DR5 spacing, indicating that the ability of RARα to activate enhancers depends on the spacing of the DRs (Fig. 6e). When we flanked the DR5 element by either an ETS binding site, which was highly enriched for RA-inducible mESC enhancers (Fig. 5d), or a SoxOct motif, we found no clear change in enhancer activity for the ETS binding site. In contrast, when the DR5 element was flanked by the SoxOct motif, we observed an increase in basal enhancer activity and also in RA-induced activation. This finding indicates a supportive role of pluripotency factors in the RA-induced enhancer activation by RARα and aligns well with the motif enrichment (Fig. 5d) and mutation analyses (Fig. 5f) for differentially active enhancers showing that the SoxOct motif is important for both basal and RA-dependent activation of RARα-occupied enhancers.

## Discussion

This study provides a comprehensive genome-wide enhancer activity map in mESCs assessed by FAIRE-STARR-seq, that can be used as a resource for further dissection of enhancer function in mESCs, and identifies various sequence features associated with enhancer activity.

To understand what discriminates active enhancers from inactive accessible regions covered by our FAIRE-STARR library, we compared these two groups and found that active regions are characterized by the presence of specific TF motifs as well as the presence of an overall higher quantity of enriched motifs (Fig. 1g). As expected, among the most prominently enriched motifs for active enhancers were the motifs of pluripotency factors OCT4, SOX2, and NANOG (cluster 18) but also SP/KLF and ETS family TFs (cluster 28 and 7), which are TFs almost ubiquitously expressed across cell types. Inactive accessible regions showed fewer enriched motifs. Moreover, these enriched motifs typically belong to cell-type specific TFs that are not expressed in mESCs and are associated with differentiation. Consistent with our findings, a recent study which systematically analyzed the quantity and composition of TF motifs for mESC enhancers described a threshold for the minimal number of TF motifs required for enhancer activity (Singh *et al*., 2021).

The introduction of UMIs during the reverse transcription step allowed us to efficiently distinguish between biological replicates and PCR duplicates of reporter-derived reads and to analyze enhancer activity quantitatively. When we started our project, UMIs were not part of the STARR-seq protocol. However, in the meantime a similar approach has been proposed for low complexity libraries where UMIs are introduced in the first PCR cycle (Neumayr *et al*., 2019). By applying UMIs, we could not only identify active enhancers, but also show that specific sequence features are associated with activity levels (Fig. 2c). For example, motifs for SP/KLF (cluster 28) and ETS TFs (cluster 7), which are essentially ubiquitously expressed across cell types, but also CG-rich motif clusters ZBTB33 (cluster 50), ZNF143 (cluster 63), ZNF263 (cluster 54), and SPIB (cluster 49) are specifically enriched at highly active enhancers and could contribute to high enhancer activity in mESCs. In contrast, motifs for pluripotency TFs (cluster 18) are similarly enriched for both highly- and lowly active enhancers and thus seem required for an enhancer to be active yet not for specifying its activity level. Thus we speculate, that individual members of the KLF and SP TF families, some of which (e.g. KLF4) were described as inhibitors of differentiation (Da and Yao, 2006; Aksoy *et al*., 2014; Tang *et al*., 2017), could cooperate with pluripotency TFs to confer high enhancer activity levels. The ETS TF family consists of many members which conduct different mainly cell-type specific and differentiation-associated functions (Sharrocks, 2001). Interestingly, ETS factor ETV5, which is very highly expressed in mESCs, was described to be essential for exit from pluripotency (Akagi *et al*., 2015; Kalkan *et al*., 2019) while ETS factor GABPA, which is the second highest expressed ETS factor in mESCs, was shown to be an activator of OCT4 (Kinoshita *et al*., 2007). Therefore, we hypothesize that specific factors of the ETS family, possibly GABPA, contribute to high enhancer activity in mESCs. On the other hand, TFs specific for differentiated cell types, such as MYOG (cluster 9), p53 (cluster 36), and POU4F1 (cluster 30) are enriched at lowly active enhancers. Since many of the TFs associated with low enhancer activity are not expressed in mESCs, we speculate that these enhancers are primed for high activity once the specific TFs are expressed *e*.*g*. at later stages during differentiation to exert their cell-type specific enhancer functions. The quantitative nature of our assay also allowed us to assess changes in enhancer activity during the early stages of RA-induced differentiation and to identify enhancers that gain or lose activity as well as associated and required TF motifs. As expected, we found that pluripotency TFs and STAT3 are associated with LIF-dependent enhancer function, however they are also found at RA-inducible promoters (Fig. 5d, S5d). Mutation experiments of individual reporter constructs (Fig. 5f) highlighted the crucial contribution of OCT4 and SOX2 motifs to enhancer activity in pluripotency, RARα::RXRα motif importance for RA-inducible activation, but also the cooperation of pluripotency and RARα::RXRα motifs in maintaining enhancer activity. Genomic deletion of selected RA-inducible enhancers (Fig. 5g, h) further validated the impact of the identified enhancers on expression of neighboring genes. Additionally, the quantitative analysis of RARα binding sites revealed a possible synergistic activation of RA-inducible RARα-bound enhancers by pluripotency TFs and RARα (Fig. 6e). A role of OCT4 in recruiting RAR::RXR to enhancers of differentiation-associated genes has been demonstrated (Simandi *et al*., 2016) and together with our data points towards an additional role of OCT4, and possibly SOX2, in facilitating increased enhancer activity during differentiation.

In our study, only a minor fraction of the probed accessible regions (4,765 of 186,959) showed significant enhancer activity in at least two out of three replicates. A recent enhancer identification study based on STARR-seq, assessing activatory potential of the whole genome, found over 18,500 active enhancers in mESCs (Peng *et al*., 2020). However, the mentioned study assessed a larger input library and did not remove PCR duplicates from their analyses, and thus applied less stringent cut-offs that would lead to similar quantities of active enhancers in our assay (*e*.*g*., we would call 21 thousand active enhancers without UMI-aware deduplication). A comparison of enhancers identified in these two studies reveals that 25.1% of the active enhancers called for our data in serum/LIF conditions without UMI-deduplicated enhancers coincide with enhancers that overlap with accessible regions from Peng *et al*. Vice versa, 33.9% of their enhancers overlap with our dataset. The difference between these studies might be related to the different promoters of the reporters that were used in these studies, since differences in promoter-enhancer compatibility can influence whether an enhancer can activate a promoter or not (Haberle *et al*., 2019). Given the different experimental set-up of these studies, these two enhancer catalogs in mESCs could complement each other.

When we analyzed the active FAIRE-STARR enhancers, which were identified by an episomal assay, for the chromatin signatures present at their endogenous genomic loci, we identified that many show the expected signature of active enhancers (high H3K4me1/H3K4me3 ratio and high H3K27ac). In addition, we identified enhancer clusters which can be classified as poised (no H3K27ac), repressed (elevated H3K27me3 or H3K9me3) or promoters (higher H3K4me3 than H3K4me1) based on their chromatin context. Strikingly, we found that almost half of the active enhancers are located at gene promoters (defined as the region up to 1 kb upstream of a TSS, Fig. 1f). The identification of such E-promoters by FAIRE-STARR-seq is in line with previous reports of promoters that act as enhancers for other genes (Dao *et al*., 2017; Diao *et al*., 2017; Dao and Spicuglia, 2018) and the high percentages of promoter-proximal enhancers identified by STARR-seq based assays in other cell types (Wang *et al*., 2018; Chaudhri *et al*., 2020). Poised enhancers (cluster F) display enrichment of TF motif cluster 24, which encompasses TFs ZIC1, ZIC3, and ZIC4. ZIC3 is a critical TF for the transition from naïve to primed pluripotency (Yang *et al*., 2019) and was shown to activate chromatin-masked enhancers in mESCs, when taken out of the endogenous context (Peng *et al*., 2020). Based on our data, we expect that that ZIC3, or another TF from the ZIC family, activates cluster F enhancers during differentiation. Furthermore, we identify a subset of enhancers (cluster G) which display enrichment of two contradictory marks: H3K36me3 associated with active transcription and H3K9me3 which marks repressed chromatin. This combination of HMs can occur on the same nucleosome (Mauser *et al*., 2017), to demark 3’ exons of zinc finger (ZNF) genes which consist of repetitive sequences (Zinc finger (ZNF) domains) (Blahnik *et al*., 2011; Hahn *et al*., 2011), and is possibly the result of two independent mechanisms, active transcription (H3K36me3) and ATRX chromatin remodeler-mediated preservation of genomic stability by repressing recombination between ZNFs (H3K9me3) (Valle-García *et al*., 2016). We now find ZNF genes to be associated with distal cluster G enhancers (Fig. 3c), which are marked with the same chromatin signature (Fig. 3a). Interestingly, the vast majority of cluster G enhancers map to endogenous retrovirus-K (ERVK) family repeats (Fig. S3f), which possess endogenous enhancer function in mESCs (Sundaram *et al*., 2017; Todd *et al*., 2019). Thus, a similar mechanism that ensures ZNF gene stability might also play a role in preventing recombination between repetitive ERVK elements that serve as enhancers of these genes.

Motivated by a study claiming that H3K122ac marks a novel class of enhancers in mESCs (Pradeepa *et al*., 2016), we added this HM to our panel of modifications assayed. However, contrary to the published study, we did not find an enhancer cluster demarked by H3K122ac while lacking the H3K27ac mark, even when we forced k-means clustering to search for more clusters (data not shown). Rather, we found that H3K122ac and H3K27ac essentially always co-occur at enhancers. This also fits with the fact that the same enzymes, histone acetyltransferases p300 and CBP, deposit both the H3K27ac and H3K122ac marks (Pasini *et al*., 2010; Tropberger *et al*., 2013). The situation is different for a subset of H3K4me3-marked promoter regions, which have high H3K122ac levels while H3K27ac levels are relatively low (Fig. S3h). Here a possible explanation is that the selective methylation of H3K27 but not H3K122 at promoter regions by polycomb repressive complex 2 prevents acetylation of H3K27 whereas H3K122 can still be modified. Accordingly, we find that H3K27me3 levels are higher at H3K27ac low, H3K122ac high regions. Taken together, these findings highlight the ability of STARR-seq to identify enhancers that are most likely poised for activation or even repressed, when taken out of the genomic context. These regions, and also promoters, are frequently excluded from enhancer prediction tools. Conversely, prediction tools like CRUP identify more putative enhancer regions that lack accessibility (Fig. S3g) but could be activated once bound by pioneering TFs. Thus, STARR-seq and enhancer prediction display complementary information about enhancers.

In summary, we generated a genome-wide enhancer activity map by FAIRE-STARR-seq which catalogs active regulatory regions in mESCs, in pluripotency and after induced differentiation. We identified features associated with enhancer activity and regulatory elements which are omitted by standard prediction tools. Our findings can serve as a reference for future functional studies of the regulatory network of genomic elements in mESCs and contribute to the refinement of computational methods to predict regulatory elements.

## Data Availability

All NGS data generated in this study was deposited at GEO (GSE171771) and published NGS data that were reanalyzed in this study are listed in Supplementary Table S2.

## Acknowledgements

We thank Sarah Kinkley and the department of computational molecular biology for insightful discussions.

