## Supplementary_Figures for "Assessing genome-wide dynamic changes in enhancer activity during early mESC differentiation by FAIRE-STARR-seq"

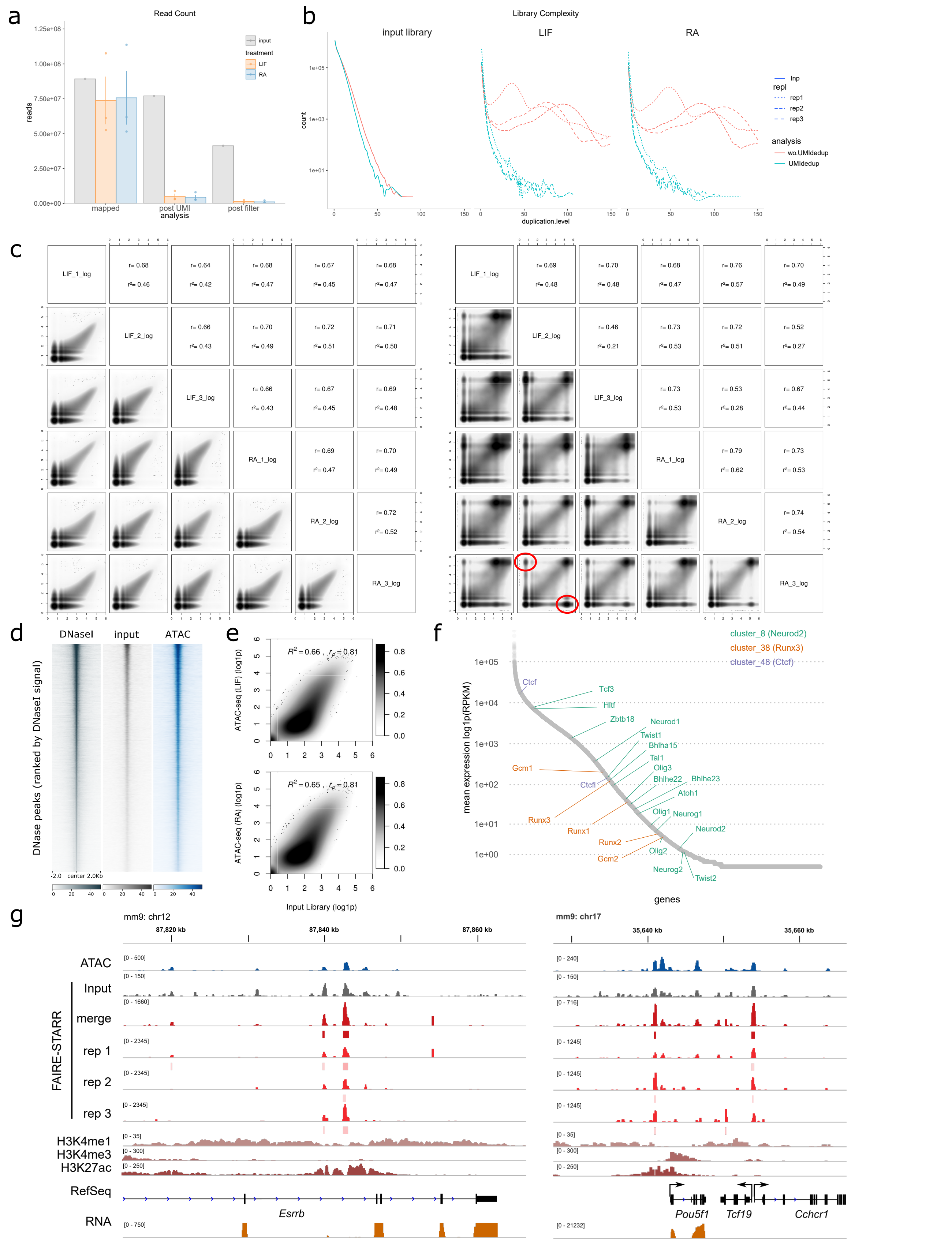


Suppl. Figure 1. FAIRE-STARR-seq in mouse embryonic stem cells.

a) Read counts of the input compared to the FAIRE-STARR-seq libraries. Absolute read numbers were counted after sequencing, after UMI-aware deduplication, and after subsequent filtering for the input and the LIF or RA treated FAIRE-STARR libraries individually. The mean of three biological FAIRE-STARR-seq replicates (bars) as well as the counts of the individual replicates (points) are shown. b) Depiction of library complexities of the input library (Inp), LIF treated, and RA treated FAIRE-STARR libraries. Data with or without UMI-aware deduplication of reads is shown for the three individual FAIRE-STARR libraries (rep1-3). c) Correlation analysis of genome-wide read distribution, comparing the three individual biological replicates for FAIRE-STARR-seq after LIF or RA treatment after (left panels) or prior to (right panels) UMI-aware read deduplication. The genome was binned into 100 bp bins and log-transformed reads per bin are plotted. Pearson correlation (r) coefficient and r-square ($r^2$) of the log-transformed data for each comparison are shown. d) Heatmaps depicting normalized read distribution of DNase-seq, FAIRE-STARR input library, and ATAC-seq at the accessible regions based on the DNase-seq data. e) Analogous to Fig. 1c, correlation analysis of genome-wide read distribution comparing the input library to ATAC-seq data of LIF- or RA-treated mESCs. Normalized and log1p transformed reads per 10 kb genomic bin are shown. f) Mean expression (RPKM normalized) for all mESC genes in pluripotency. TF genes belonging to motif clusters 8, 28, and 48 are highlighted. g) IGV browser view of exemplary genomic regions illustrating the STARR activity from three individual biological replicates (rep1-3) and the merged signal. RPGC normalized signals for FAIRE-STARR replicates and input are shown.

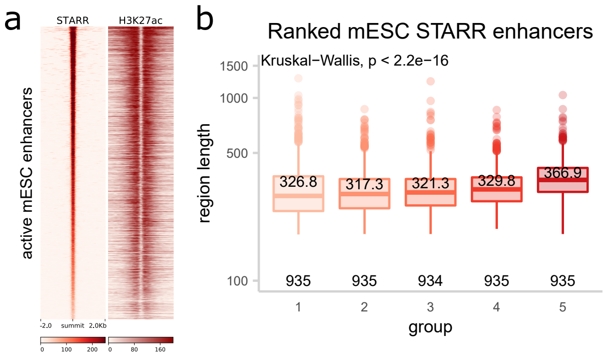


Suppl. Figure 2. FAIRE-STARR-seq enables quantification of enhancer activity and activity level-associated sequence features.

a) Normalized FAIRE-STARR and H3K27ac signal distribution at FAIRE-STARR enhancers ranked by STARR activity. b) Average sequence length for each group of FAIRE-STARR enhancers (grouping as depicted in Fig. 2a). Boxplots depict the length-distribution of all genes per group, whiskers extend to 1.5 IQR. P-values were calculated by Kruskal-Wallis test for differences between all groups and group sizes are indicated directly above the X-axis.

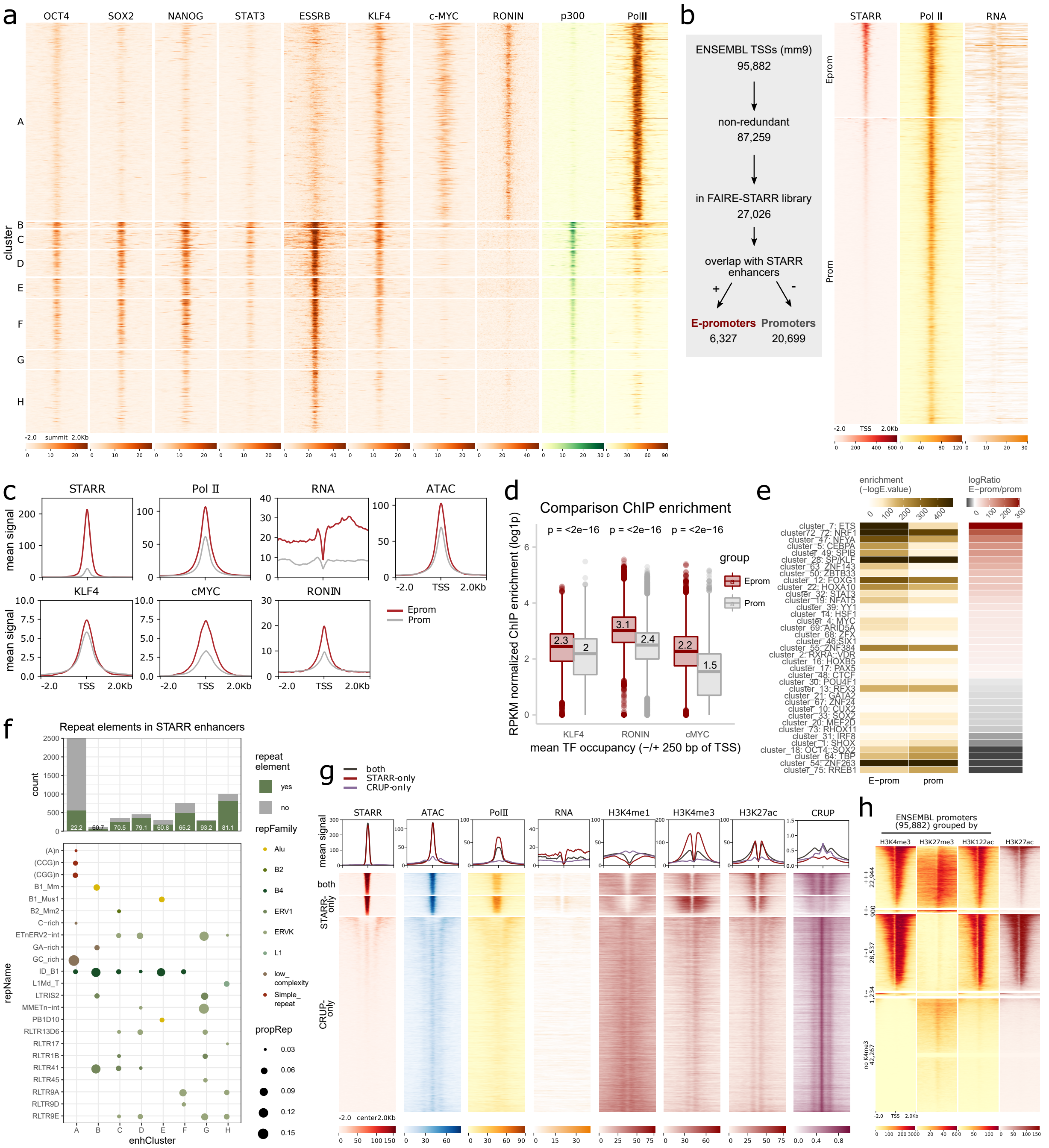


Suppl. Figure 3. Functional mESC enhancers reside in different epigenomic environments.

a) Distribution of TFs as indicated, p300, and PolII at active FAIRE-STARR enhancers, clustered as depicted in Fig. 3a. b) TSSs (ENSEMBL annotation mm9) were filtered for redundancy and coverage in the FAIRE-STARR library and subsequently divided into those which do (E-promoters, Eprom) or do not (regular promoters, Prom) overlap with active FAIRE-STARR enhancers. The two groups were ranked by their STARR signal and PolII and RNA enrichment at these regions was plotted. c) Anchor plots showing the mean enrichment of signals as indicated at E-promoters and regular promoters (grouped as in b). d) Comparison of TF enrichment at E-promoters and promoters (-/+ 250 bp from TSSs). Boxplots depict the distribution of RPKM normalized ChIP signal (log1p transformed data) and p-values for unpaired Wilcoxon tests comparing enrichment for E-promoters and promoters. e) TF motif enrichment for E-promoters and (6327 randomly selected) promoters was performed with AME using the JASPAR 2018 clustered vertebrate motif database. Significance of enrichment (-log10E <= 1e-4) for differentially enriched (-log ratio >= 5) motifs are shown. f) Intersection of FAIRE-STARR enhancer clusters with elements from RepeatMasker (Smit AFA, Hubley R, 2013) for the mm9 genome. The bar plot shows the absolute count for repeat elements per enhancer group and the percentage of all repeats per group are indicated. The lower panel shows the proportion of individual repeats, color-coded by repeat family, for each cluster. Only repeats which make up at least 3% of all repeats per cluster are shown. g) Mean enrichment (upper panels, anchor plots) and distribution (heatmaps) of STARR-, ATAC-, RNA-, selected HM ChIP-seq signals, as well as enhancer probability by CRUP prediction for enhancers identified only by FAIRE-STARR, only by CRUP or by both. h) Distribution of selected HMs at ENSEMBL promoters (95,882), which were grouped by overlap with significant enrichment (by peak calling) of H3K4me3, H3K27me3, and H3K122ac. +++: H3K4me3, K3K27me3, and H3K122ac positive. ++-: H3K4me3, K3K27me3, but no H3K122ac. +-+: H3K4me3 and H3K122ac positive, but no K3K27me3. +--: H3K4me3 positive, but no K3K27me3 or H3K122ac. No H3K4me3: Promoters without significant H3K4me3 enrichment. H3K27ac enrichment was plotted for the promoters grouped as described above.

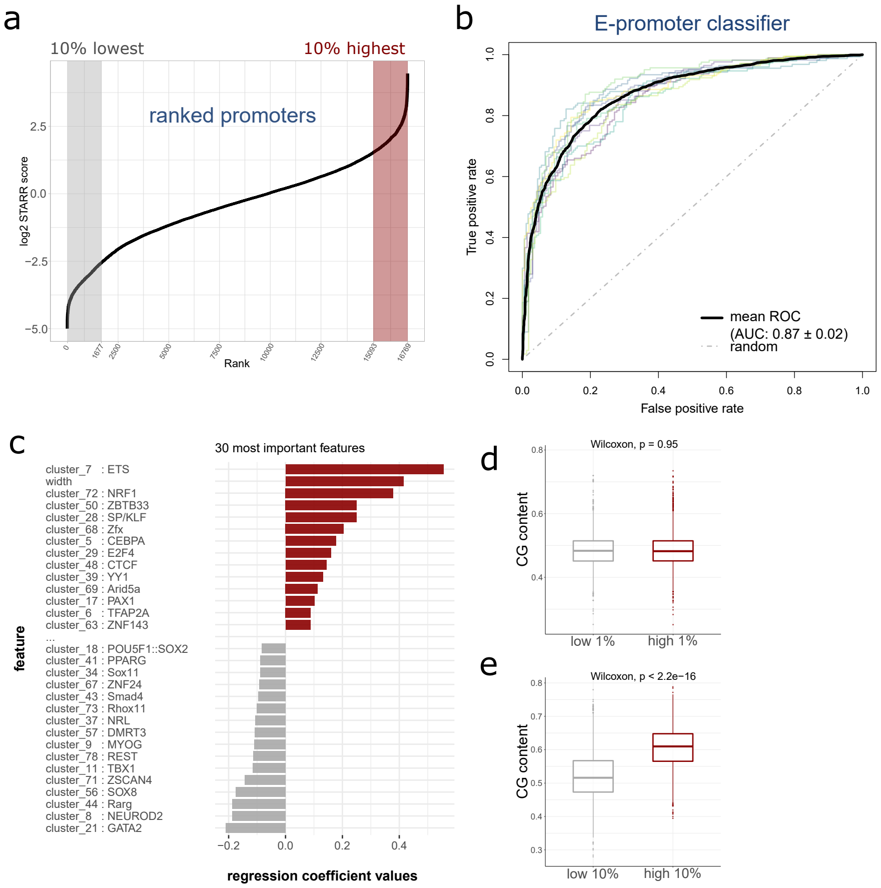


Suppl. Figure 4: Sequence-based prediction of enhancers and E-promoters.

a) ENSEMBL promoters overlapping with FAIRE-STARR library regions (Fig. 4a) were ranked for their STARR score and the 10% highest and lowest ranking promoters were used for model building. b) Receiver operating characteristic (ROC) curve for E-promoter prediction model performance for each of the outer cross-validation folds and mean and standard deviation of the area under the ROC curve are shown. c) The 30 most predictive variables for the optimal model of E-promoter prediction and their coefficients are shown. Positive coefficients indicate a positive association with high STARR scores, while motifs with negative coefficients are associated with low-scoring elements. Comparison of CG content of high- and low-ranking d) enhancers and e) E-promoters. P-values for Wilcoxon test are depicted.

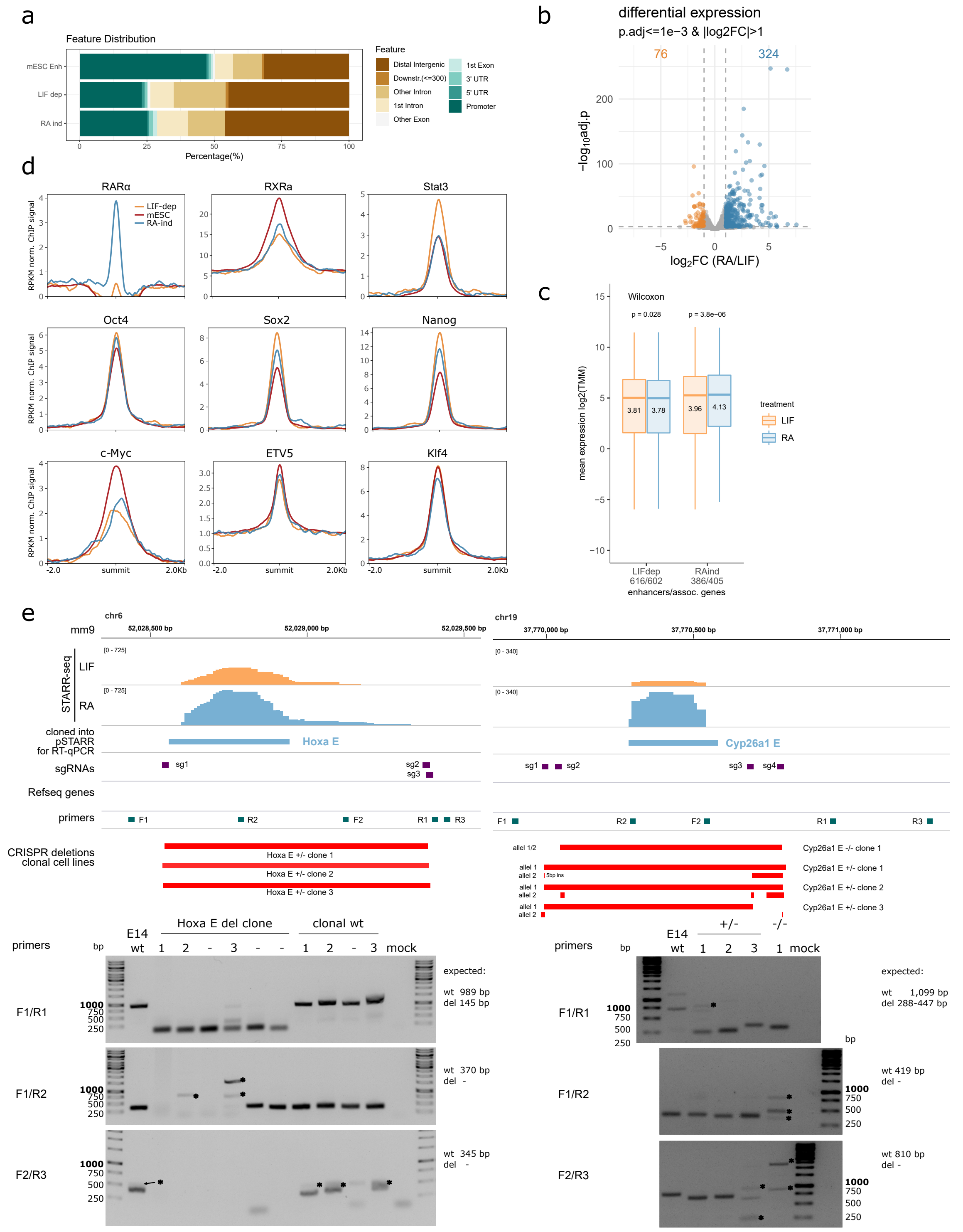


Suppl. Figure 5: Differentiation-associated changes in enhancer activity.

a) Genomic distribution of active enhancers in mESCs, of LIF-dependent and RA-inducible enhancers with respect to annotated Refseq genes. Promoters were defined as the regions 1 kb upstream of a TSS. b) Differential gene expression comparing E14 cells treated for 4 h with either LIF or RA using DESeq2. Significantly up- (blue) and down-regulated (orange) genes and cut-offs are indicated. c) Genes were paired with enhancers by distance using GREAT and TMM-normalized gene expression counts per enhancer group and treatment are shown. P-values from paired Wilcoxon tests are shown. d) Mean normalized enrichment of TFs as indicated at LIF-dependent, RA-inducible, and active mESC STARR enhancers. e) Genotyping of E14 enhancer deletion CRISPR/Cas9 clones. Upper panels depict the targeted genomic regions (mm9) and FAIRE-STARR-seq signals (LIF or RA treated). Genomic locations of regions cloned into pSTARR for RT-qPCR (blue, Fig. 5e and f), guide RNAs for targeting Cas9 (sgRNAs, purple), primers (green) used for genotyping PCRs, and detected deletions of the individual clones are depicted. Lower panels show genotyping PCR results for genomic DNA recovered from individual clones or parental line (E14 wt) using primer pairs as indicated (locations shown in the upper panel). Asterix mark unspecific PCR bands. Expected PCR amplicon sizes for deletion (del) or wild type (wt) clones are indicated.

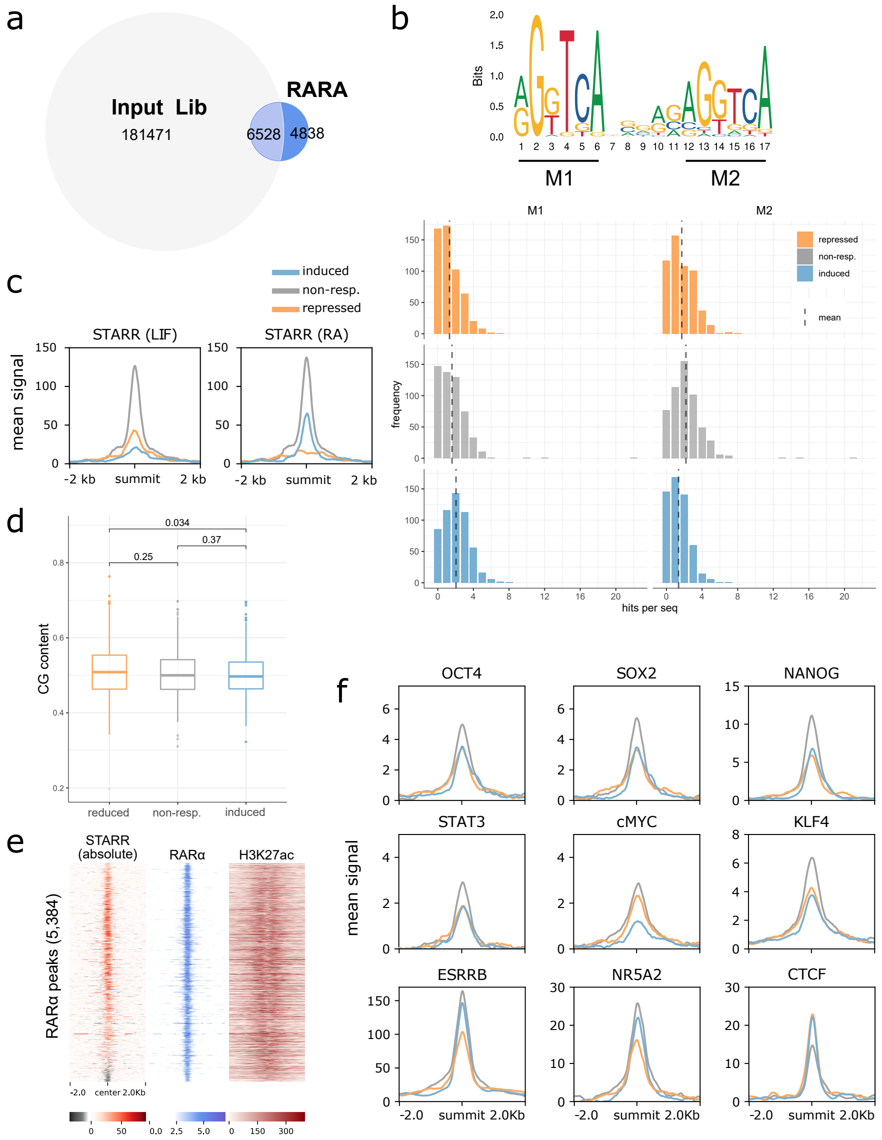


Suppl. Figure 6: Identification of enhancer activity-associated features of RARα binding sites.

a) Intersection of FAIRE-STARR-seq input library and RARα-occupied sites. 6,528 RARα sites covered by our FAIRE-STARR-seq input library were used for further analysis. b) Frequencies of the RARα::RXRα consensus motif (MA0159.1) repeat half-sites (M1 and M2, upper panel) at repressed, non-responsive, and induced RARα-occupied sites. c) Absolute FAIRE-STARR activity at repressed, non-responsive, and induced RARα-occupied sites in LIF or RA treated cells. d) CG content distribution at repressed, non-responsive, and induced RARα-occupied sites. P-values were derived from Wilcoxon tests. e) Heatmap representing distribution of FAIRE-STARR signal, RARα and H3K27ac enrichment at RARα-occupied sites which were ranked by their logSTARR score (RA/LIF). f) Average enrichment of selected TFs at induced (blue), non-responding (gray), or repressed (orange) RARα-occupied sites. All ChIP-seq data were derived from pluripotent mESCs without RA treatment.
